# Neural ensembles for music production recruit more language instruments as rhythmic complexity increases

**DOI:** 10.1101/2025.11.16.688175

**Authors:** Ut Meng Lei, Yan Tian, Mengxiao Wang, Yihong Mu, Chi Un Choi, Katy Ieong Cheng Ho Weatherly, Martin I. Sereno, Defeng Li, Victoria Lai Cheng Lei, Ruey-Song Huang

**Author notes:** **Corresponding Authors:** Ruey-Song Huang, Victoria Lai Cheng Lei **Email:**.

## Abstract

Language and music, two core forms of human communication, both rely on rhythmic control. To date, the neural ensembles supporting their real-time production are not fully characterized. Using rapid phase-encoded fMRI, we mapped the spatiotemporal brain dynamics of overt language and music production under varying rhythm control demands. We observed overlapping hemodynamic traveling waves across domains, with auditory-motor regions activated earlier, followed by visual, posterior parietal, and Sylvian parietal-temporal (Spt) regions, supporting sensorimotor transformation and online monitoring of production. Increasing rhythmic complexity elicited slower and stronger activations in both domains: compared with baseline, regular rhythm control elicited delayed and amplified responses, while irregular rhythm control produced the slowest and strongest responses. During music production, rhythm-related activity expanded into frontal and temporal language regions, suggesting that higher rhythmic demands engage language resources. These findings reveal dynamic neural ensembles that can flexibly reconfigure shared resources to support temporal precision in sequence production.

## Introduction

Rhythmic control, the precise timing and coordination of motor, auditory, and cognitive processes, is central to both language and music production, two of the most distinctive human capacities. Both systems depend on hierarchical timing and on real-time sequencing and motor planning that coordinate multimodal brain regions^1,2^. These shared demands on timing and rhythm have fueled interest in common neural resources, yet the neural dynamics during overt rhythmic production of language and music remain underexplored with noninvasive neuroimaging. Meter, a hierarchical pattern of strong and weak beats, organizes temporal prediction and enables listeners to anticipate salient events (stressed syllables or accented notes)^2-6^. By aligning internal timing and motor planning to a metric pulse, meter supports predictive sequencing and chunking of elements. Examining how metric structure shapes anticipatory motor commands and sensory processing during overt production can therefore reveal the mechanisms the brain uses to encode, plan and execute structured sequences across language and music.

Theories and models have been proposed to delineate the interaction and coordination between different brain regions during language and music rhythm processing. For example, Rauschecker^7^ proposed a model to show the dynamic interplay between auditory and motor systems during sensorimotor integration, where efference copies from motor regions generate predictions for sensory input. The model of Action Simulation for Auditory Prediction (ASAP)^8^ builds directly on this architecture, proposing two-way communication between auditory and motor planning regions in the prediction of rhythm, where motor planning regions simulate periodic movements and send predictive signals to auditory regions, thereby facilitating the anticipation of upcoming beats. Likewise, the Processing Rhythm in Speech and Music (PRISM)^9^ framework describes auditory-motor coupling as supporting timing prediction in music and speech. These frameworks align with the dual-stream models of the auditory cortex, originally proposed for speech^10,11^ and later extended to music processing^12^, outlining a dorsal stream that supports sensorimotor integration and predictive timing, and a ventral stream involved in sound recognition and auditory object identification.

Previous functional magnetic resonance imaging (fMRI) studies have identified a bilateral auditory-motor network consisting of frontal, parietal, and temporal regions that are associated with musical rhythm processing^13-30^. Within this network, bilateral supplementary motor area (SMA), pre-SMA, dorsal premotor cortex, prefrontal cortex, inferior parietal lobe (IPL), and right superior temporal regions were found to be modulated by rhythmic complexities. Moreover, some studies suggest that musical rhythm processing engages brain regions implicated in higher-order language processing, including the Broca’s area, planum temporale, left supramarginal gyrus (SMG), and insula^20,21,29,30^. Other studies examining neural correlates of language rhythm processing have likewise highlighted the involvement of SMA, cingulate gyrus, SMG, superior temporal gyrus (STG), inferior frontal gyrus (IFG), and insula^31-33^. However, direct comparisons of the neural mechanisms between language and music rhythm processing remain scarce^2^, especially for overt production of comparable sequences, such as speech versus wind instrument playing – both involve orofacial, vocal tract, and respiratory control. However, achieving naturalistic and continuous overt production of language and music sequences with real-time auditory feedback has been challenging for fMRI, given its sensitivity to head motion and to interference from scanner noise. As a result, most prior work has relied on covert tasks or constrained production paradigms and has emphasized static activation contrasts over dynamic interactions among brain regions.

Consequently, although the neural correlates of language and music processing in general have been extensively examined^13-30,34-39^, the spatiotemporal brain dynamics remain largely unexplored with fMRI. Some studies have revealed functional connections between auditory and motor regions during rhythm processing, but the directional flow of signals and the dynamics of their interactions are yet to be fully investigated^8^. Existing models shedding light on information flows between brain regions during language and music rhythm processing thus remain primarily conceptual. To what extent overt language and music production recruit overlapping brain regions and pathways and how crossmodal interactions unfold over time and under conditions with varying rhythmic complexities remain open questions.

To map brain regions involved in rhythm processing, some fMRI studies have investigated various effectors (mouth, fingers, and legs) linked to language and music production, and identified regions involved in the encoding, maintenance, production, and synchronization stages of rhythm production^24,25,27^. For example, the STG, IFG, and IPL encode rhythm information from auditory input, whereas during rhythm retrieval, the IPL and SMA transformed temporal representations of hierarchical sequences in the fronto-parietal network into motor sequences that drive output. However, due to the inherent constraints of contrast-based fMRI designs and analyses, these studies are restricted to locating brain regions engaged in discrete phases of rhythm processing, rather than tracking the continuous flow of brain activities throughout the encoding-to-production process.

Addressing the existing gaps, this study applied rapid phase-encoded fMRI^40,41^ to capture the whole-brain spatiotemporal dynamics while overcoming head-motion challenges associated with overt production tasks in both domains. Language and music sequences imposing varying rhythm control demands were used as stimuli during naturalistic speech production and wind instrument playing by amateur musicians, enabling ecologically valid production tasks in both domains. Rhythm control demands were operationalized as the degree of regularity required in producing alternating strong and weak elements. Within each domain, three conditions were implemented: A self-paced baseline condition requiring no explicit rhythm control, and congruent and incongruent conditions requiring the production of sequences with prompted regular and irregular accents, respectively. By providing direct neuroimaging evidence of how brain activations unfold across regions during the real-time perception-to-production process, this study delineates and compares the spatiotemporal patterns of information flows under different rhythmic conditions, which substantiate and refine existing models about the neural pathways underlying language and music processing.

## Results

Thirty native Cantonese speakers were scanned while participating in six periodic event-related tasks involving the overt production of English sentences or musical phrases (Fig. 1 and Methods). For each task, Fourier-based analysis of hemodynamic time series revealed strong periodic hemodynamic signals at 16 cycles per scan, which were displayed with color-coded activation phases on the cortical surface (Fig. 1d).

**Fig. 1.**
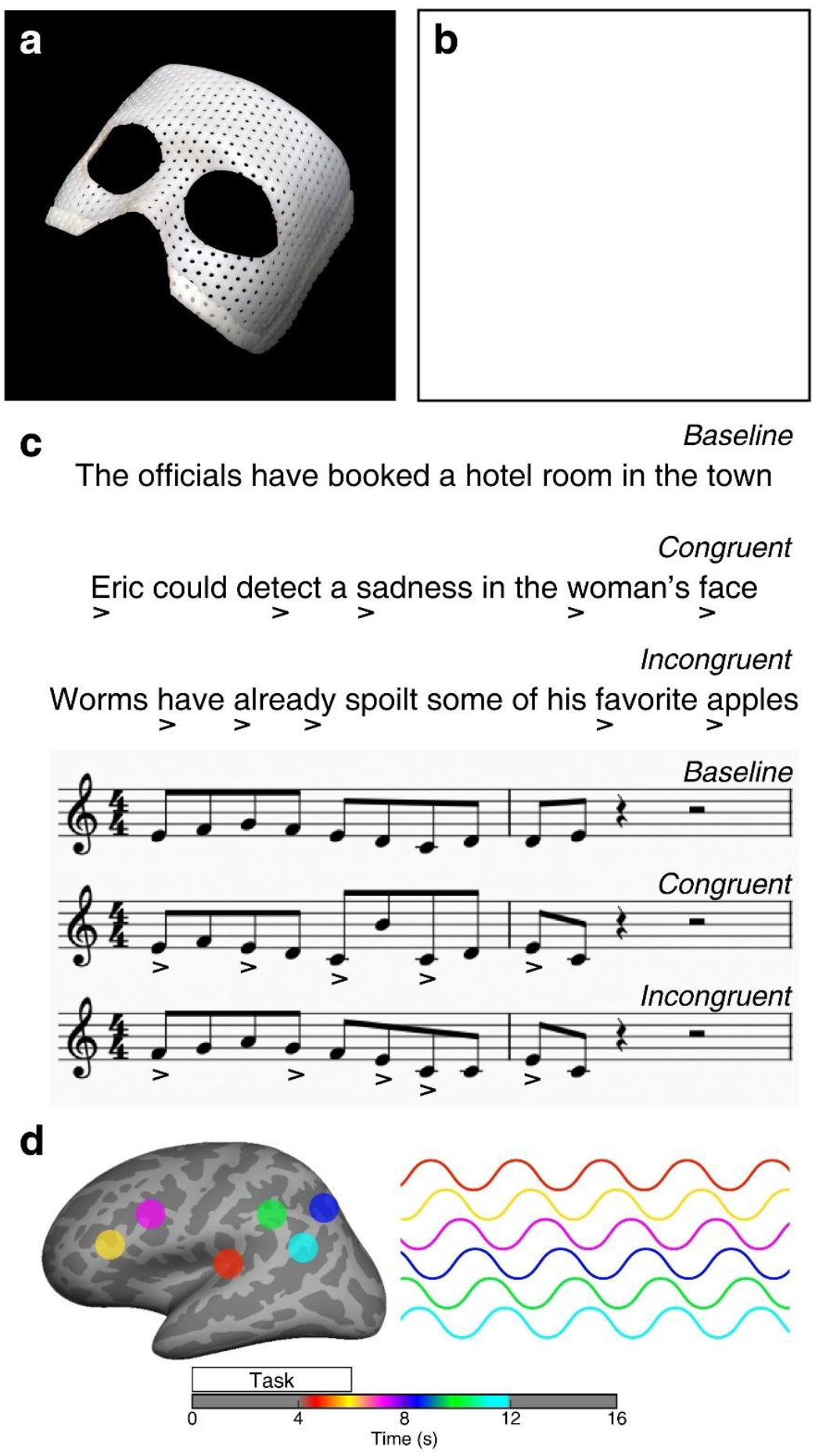
Experimental setup and design. **a**, A custom-molded mask prevents head motion during overt language and music production. **b**, Experimental setup for recorder playing tasks. **c**, Sample stimuli for language or music tasks with baseline (without accent symbols), congruent (with regular accent symbols), and incongruent (with irregular accent symbols) conditions. **d**, A schematic neural ensemble map illustrating phase-encoded activations in the left hemisphere. Periodic hemodynamic signals in six cortical regions are color-coded by their activation phases, as indicated by the colorbar.

### Spatiotemporal unfolding of phase-encoded activations during language and music production

Group-average neural ensemble maps show the overall extent of statistically significant phase-encoded activations (*n* = 30, *F*(2, 58) = 4.99, *P* < 0.01, cluster corrected) for each language or music task (Figs. 2, 3 and Methods). Color-coded brain regions indicate their activation at different timings (phases), collectively functioning like a coordinated neural ensemble to perform each task. All language and music tasks commonly activated bilateral primary visual cortex (PVC), early visual cortex (EVC), dorsal stream visual cortex (DSVC), ventral stream visual cortex (VSVC), MT+ complex neighboring visual areas (MT+CNVA), somatosensory and motor cortex (SMC), paracentral lobular and midcingulate cortex (PLMC; including SMA), premotor cortex (PMC), posterior opercular cortex (POC), early auditory cortex (EAC), auditory association cortex (AAC), insular and frontal opercular cortex (IFOC), lateral temporal cortex (LTC), temporal-parietal-occipital junction (TPOJ), superior parietal cortex (SPC), inferior parietal cortex (IPC), posterior cingulate cortex (PCC), anterior cingulate and medial prefrontal cortex (ACMPC), inferior frontal cortex (IFC), and dorsolateral prefrontal cortex (DLPFC) (Figs. 2, 3 and Supplementary Table 1). Furthermore, recorder playing in music tasks activated bilateral manual control regions in SMC, PLMC, PMC, IPC, SPC, and POC (Figs. 2b, 3a).

**Table 1.** Percentages of overlapping and domain-specific vertices in each task.

|  | Left hemisphere |  |  |  | Right hemisphere |  |  |  |
| --- | --- | --- | --- | --- | --- | --- | --- | --- |
|  | Language |  | Music |  | Language |  | Music |  |
| Baseline | 29.12% | 70.88% | 75.88% | 24.12% | 29.44% | 70.56% | 69.55% | 30.45% |
| Congruent | 15.34% | 84.66% | 74.12% | 25.88% | 11.50% | 88.50% | 68.97% | 31.03% |
| Incongruent | 14.10% | 85.90% | 74.32% | 25.68% | 12.46% | 87.54% | 71.74% | 28.26% |

Note: The value in each white cell indicates the percentage of vertices significantly activated by each task alone (Fig. 3a, columns 1 and 2). Values in shaded cells indicate the percentage of vertices significantly activated by both language and music tasks, corresponding to the cyan overlap regions shown in the conjunction maps (Fig. 3a, column 3).

**Fig. 2.**
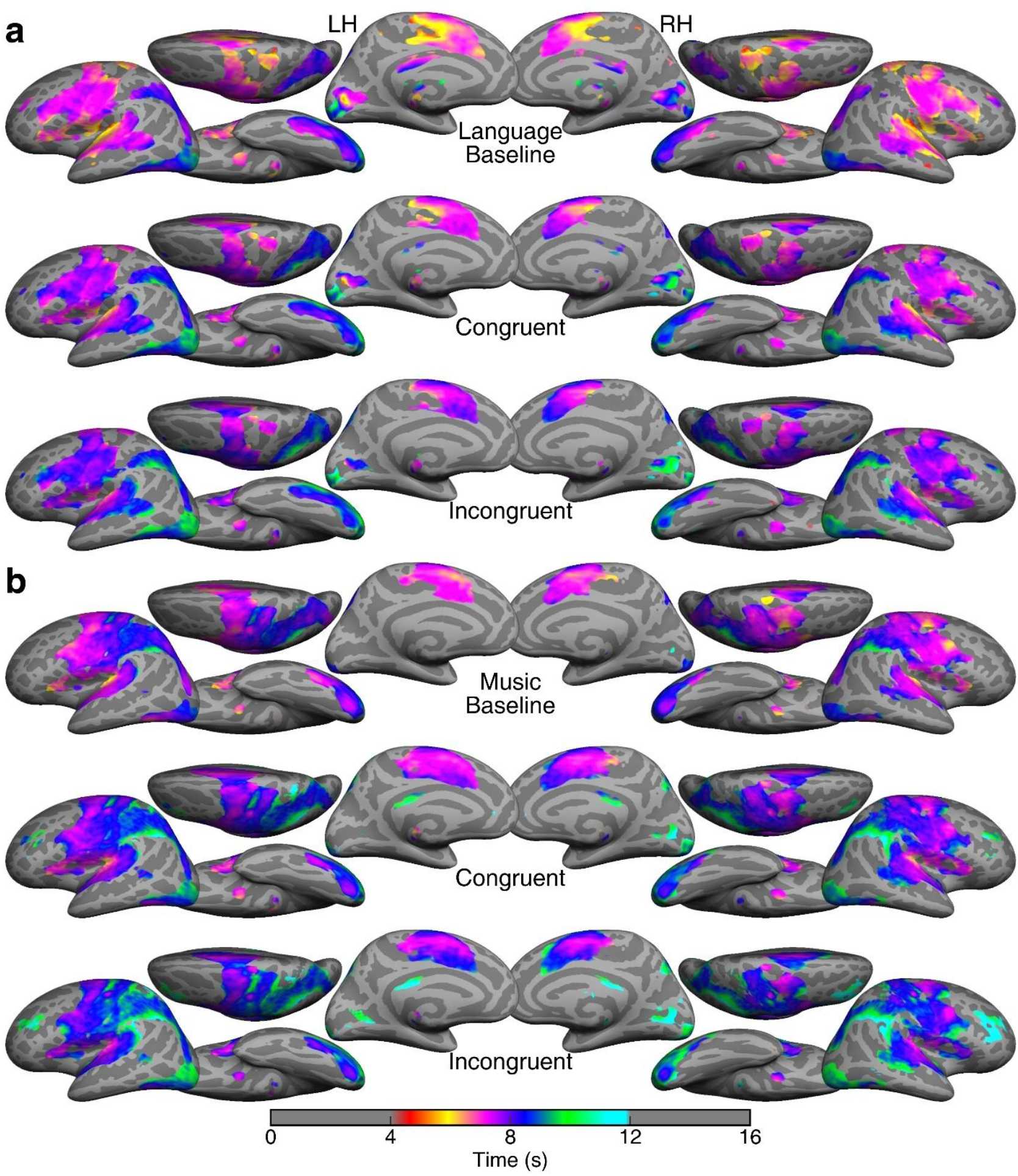
Group-average neural ensemble maps shown on inflated cortical surfaces. **a**, Phase-encoded activations in three language tasks. **b**, Phase-encoded activations in three music tasks. Statistically significant periodic hemodynamic signals (16 cycles/scan; *n* = 30, *F*(2, 58) = 4.99, *P* < 0.01, cluster corrected) are displayed in colors according to their phases (time delays). Only task-related activations between 4 and 12 s (due to hemodynamic response delay) are shown. LH: left hemisphere; RH: right hemisphere.

**Fig. 3.**
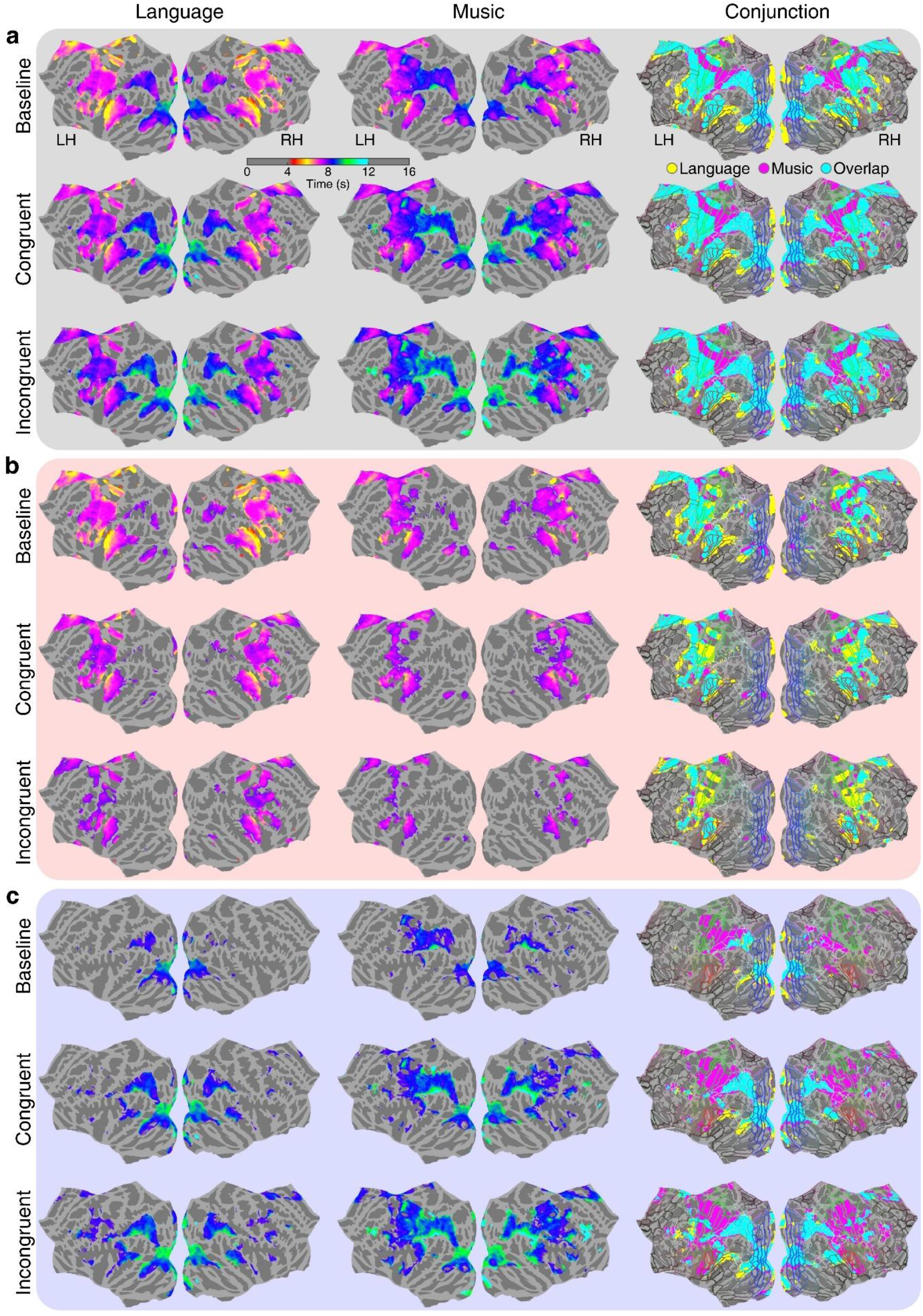
Contrasting phase-specific activation patterns between language and music tasks. Maps of statistically significant activations (*n* = 30, *F*(2, 58) = 4.99, *P* < 0.01, cluster corrected) in Fig. 2 are redisplayed on flattened cortical surfaces for **a**, full-range (4-12 s); **b**, earlier phases (4-8 s); and **c**, later phases (8-12 s). The third column in each panel shows the surface-based conjunction of activation maps from the first two columns, overlaid with HCP-MMP1 parcellation (Supplementary Fig. 1 and Methods). Yellow: regions activated by language tasks only; Magenta: regions activated by music tasks only; Cyan: regions activated by both language and music tasks. LH: left hemisphere; RH: right hemisphere.

The continuous process of each language or music production task unfolds as sequential activations across the cortical surface, with their phases represented by different colors (see animation of traveling waves in Supplementary Movies 1-6). For example, in the language congruent task (Figs. 2a, 3a), the left temporal lobe shows a progression of activation phases traveling from EAC to AAC and finally reaching the Sylvian parietal-temporal area^38,42^ (Spt) at the posterior lateral sulcus, which is visualized as a continuous color gradient transitioning from yellowish to pinkish, bluish, and greenish hues.

While the overall activation extent remains relatively stable across tasks within each domain, the activation phases show increasing delays from baseline through congruent and incongruent conditions, as indicated by the expansion of bluish-greenish areas in the neural ensemble maps (Figs. 2, 3). This progressive shift in activation phases demonstrates consistent modulation of rhythm control demands in both domains, as detailed in the following sections.

### Rhythm control demands shift spatiotemporal activation patterns in both domains

To compare the spatiotemporal activation patterns across tasks and between domains, we displayed phase-encoded activations and overlaps on flattened cortical surfaces for each condition (Fig. 3). The conjunction maps between baseline language and music tasks (Fig. 3a, column 3; Table 1; Supplementary Table 1c) reveal large overlaps in the dorsal and ventral visual streams (DSVC and VSVC), posterior parietal cortex (SPC and IPC), early and association auditory cortex (EAC and AAC), anterior insular, as well as supplementary motor, premotor, and primary sensorimotor cortex (PLMC, PMC, and SMC). In particular, both overt speech production and recorder playing activated the orofacial (inferior portion) and respiratory control (superior portion) regions in SMC, which are clearly separated by the hand representations in SMC (magenta region in the conjunction maps). Furthermore, music tasks uniquely activated manual control regions in premotor, posterior parietal, secondary somatosensory, and posterior cingulate cortex. Beyond the regions commonly activated by the music baseline task, the language baseline task activated more extended regions in the bilateral DLPFC, IFC, superior temporal sulcus (STS), larynx region in opercular cortex, left LTC, as well as the peripheral visual field representations in primary visual cortex (V1).

The overlaps between language and music tasks under explicit rhythmic conditions, whether with congruent or incongruent meter, were more extensive than those in the baseline (Fig. 3a, column 3). The expanded overlaps were found in DLPFC, IFC, IFOC, EAC, ACC, TPOJ, POC, PLMC, IPC, SPC, and PCC. When quantified from the language domain, the proportion of vertices commonly activated by music tasks increased substantially across conditions in the left hemisphere (Table 1). This trend suggests that, as rhythmic control demands increase, music-related activities progressively encroach into language-dominant cortical areas. In contrast, the proportion of music-specific activations remained relatively stable, indicating regions involved in manual control during recorder playing.

To determine how increasing rhythm control demands modulate the spatiotemporal activation patterns, we divided each activation map into earlier-phase regions (4-8 s; Fig. 3b) and later-phase regions (8-12 s; Fig. 3c) (Supplementary Table 1b). These two ranges were selected empirically to facilitate subsequent analyses and interpretations. Regions with leading activations could be involved in preparatory processes such as production initialization and sequence planning, whereas the regions activated subsequently could be involved in the execution processes underlying continuous rhythm production and online monitoring. The changes in the spatial extent of activations across conditions reflect temporal delays in processing along with increasing rhythm control demands. For instance, the language and music baseline tasks showed broader earlier-phase activations and more limited later-phase activations, indicating faster processing in both domains. Relative to baseline conditions, in the incongruent conditions, the regions with earlier-phase activations shrank while the regions with later-phase activations expanded, suggesting delayed processing with more activations falling into the later phases.

In earlier phases, substantial overlaps were primarily found in SMC, PMC, PLMC (including SMA), EAC, SPC, and IPC (Fig. 3b, column 3), with the language baseline condition exhibiting slightly more extensive activation in these regions. The reduced activation in the congruent and incongruent conditions compared to baseline conditions for both domains suggests a progressively delayed response (initialization) as rhythm control demands increase. The overlapping regions retained during the earlier phases of the incongruent conditions indicate the essential leading regions that initiate rhythm production in both domains (Fig. 3b, column 3, cyan regions).

In later phases, overlaps were mainly found in the visual and posterior parietal cortices, which were less extensive than the overlaps in earlier phases (Fig. 3c, column 3). Music tasks additionally activated manual control regions in the later phases. Together, increasing rhythm control demands shift spatiotemporal activation patterns in both domains by delaying brain responses across conditions, as reflected by shrinking regions with earlier-phase activations and expanding regions with later-phase activations.

### Music production recruits language-dominant regions with increasing rhythm control demands

To determine how regional brain activations were modulated by different rhythm control demands, we compared the changes in vertex-wise activation ratios and phase distributions within each surface-based region of interest (sROI) based on the HCP-MMP1 parcellation (Supplementary Figs. 1-4, Supplementary Table 1a, 1b, and Methods). Four types of changes in spatiotemporal patterns across conditions in selected sROIs in the left hemisphere are illustrated in Fig. 4.

**Fig. 4.**
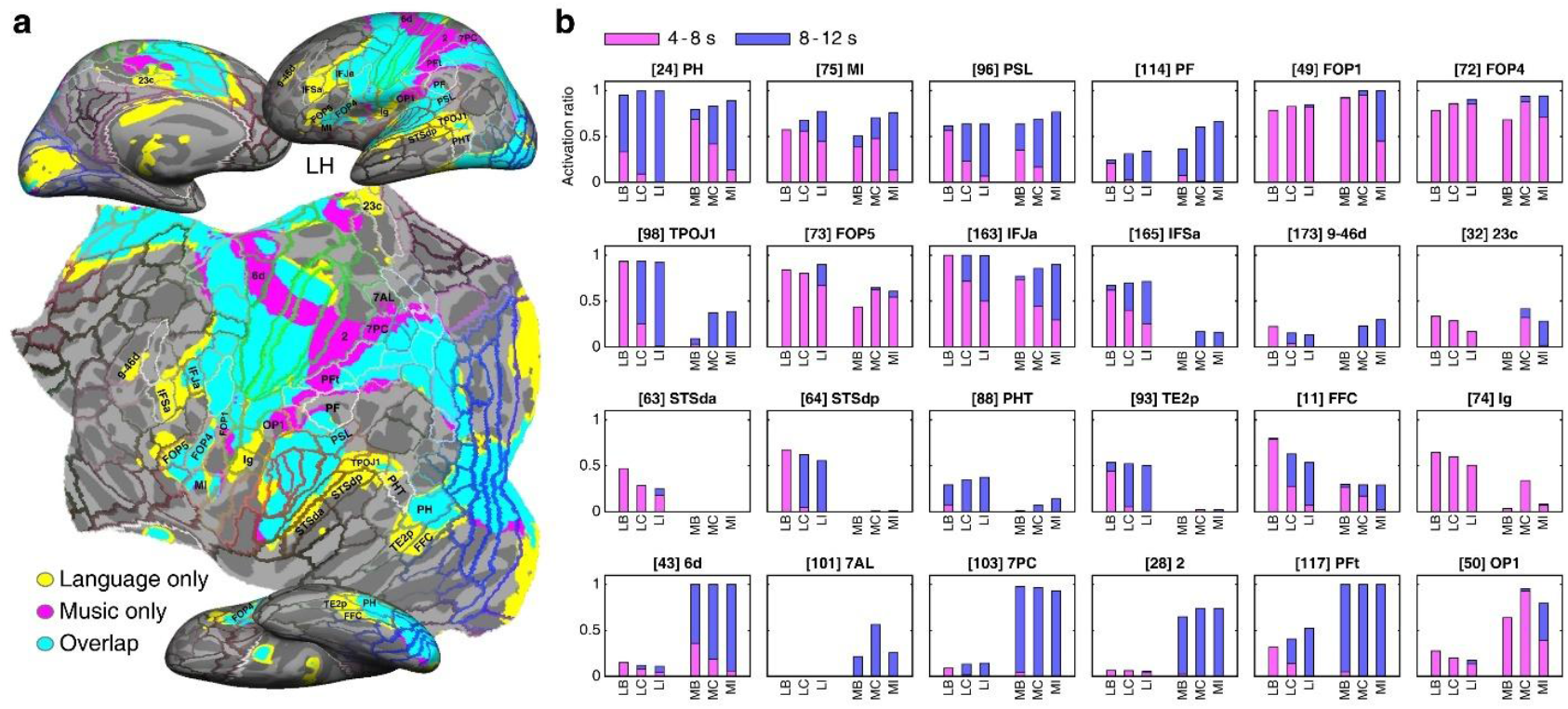
Contrasting activation phases across tasks and between domains within surface-based regions of interest. a., The left-hemisphere conjunction map between baseline language and music tasks (Fig. 3a, column 3) is redisplayed here on inflated and flattened cortical surfaces, overlaid with HCP-MMP1 parcellation. **b**., Activation ratios and phase distributions across conditions within each of the selected sROIs labeled in Fig. 4a. The number in the square brackets indicates the sROI index (see Supplementary Table 1b). The height of each stacked bar indicates the overall ratio of vertices with full-range (4-12 s) activations in each sROI. Pink and blue segments represent the ratios of vertices with earlier-phase (4-8 s) and later-phase (8-12 s) activations, respectively. LB, LC, and LI: baseline, congruent, and incongruent conditions in language tasks; MB, MC, and MI: baseline, congruent, and incongruent conditions in music tasks.

First, a trend of gradual increase in the activation strength and time delay was observed across conditions in both domains. The overall activation ratios in areas PH, MI, PSL, PF, FOP1, and FOP4 slightly increased (as indicated by longer stacked bars) with rhythm control demands in both domains (Fig. 4b, row 1). At the same statistical threshold, the increased ratios also reflect higher activation amplitudes in congruent and incongruent conditions. Furthermore, the expansion of later-phase (blue) segments indicates activation delays, reflecting increasing demands in rhythm control from baseline through congruent to incongruent conditions. Similar modulations of activation strength and phase can be observed in the corresponding sROIs in the right hemisphere (Supplementary Figs. 2–4).

Second, several language-dominant regions in STS and DLPFC were more engaged in music tasks under increasing rhythm control demands. Areas TPOJ1, FOP5, IFJa, IFSa, 9-46d, and 23c were primarily activated by language tasks, with slight variation in activation ratios across language conditions (Fig. 4b, row 2). These areas exhibited increased activations in music congruent and incongruent conditions, compared with the baseline. A similar pattern can be observed in the corresponding sROI in the right hemisphere (Supplementary Figs. 2-4).

Third, areas STSda, STSdp, PHT, TE2p, FFC, and Ig were primarily activated in all three language conditions (Fig. 4b, row 3). Among them, PHT and Ig were partially activated in music congruent and incongruent conditions. While these regions remained primarily language-dominant in the right hemisphere, areas PHT and FFC in the ventral occipitotemporal cortex and STSdp additionally exhibited expanded music-related activations under conditions requiring greater rhythm control (Supplementary Figs. 2–4).

Fourth, areas 6d, 7AL, 7PC, 2, PFt, and OP1 in the left hemisphere displayed music-related activations with limited to no response to language tasks (Fig. 4b, row 4). These areas, located in premotor, sensorimotor, secondary somatosensory, and posterior parietal cortex, were involved in manual control in all three music conditions, and were partially activated in all three language conditions, except 7AL. A similar pattern was observed in the corresponding sROIs in the right hemisphere (Supplementary Figs. 2-4).

While both domains exhibited bilateral activations, they were left-dominant, as indicated by the laterality index (LI) maps (Supplementary Fig. 1c). Moreover, the expansion of music-related activations under higher rhythm control demands was bilateral, with more pronounced increases in language-dominant regions including STS (TPOJ1), DLPFC and IFC (9-46d, IFSa, and IFJa), and VSVC and LTC (PHT and FFC).

### Rhythm control demands modulated the amplitude, phase, and spatial distribution of activations

To compare the overall and fine-grained spatiotemporal activation patterns across conditions, we computed a surge profile depicting the distribution of activation phases within each hemisphere as well as a hemodynamic Gantt chart comprising the surge profiles (horizontal bars) in 180 sROIs per hemisphere based on the HCP-MMP1 parcellation (Fig. 5, Supplementary Fig. 1, and Methods).

**Fig. 5.**
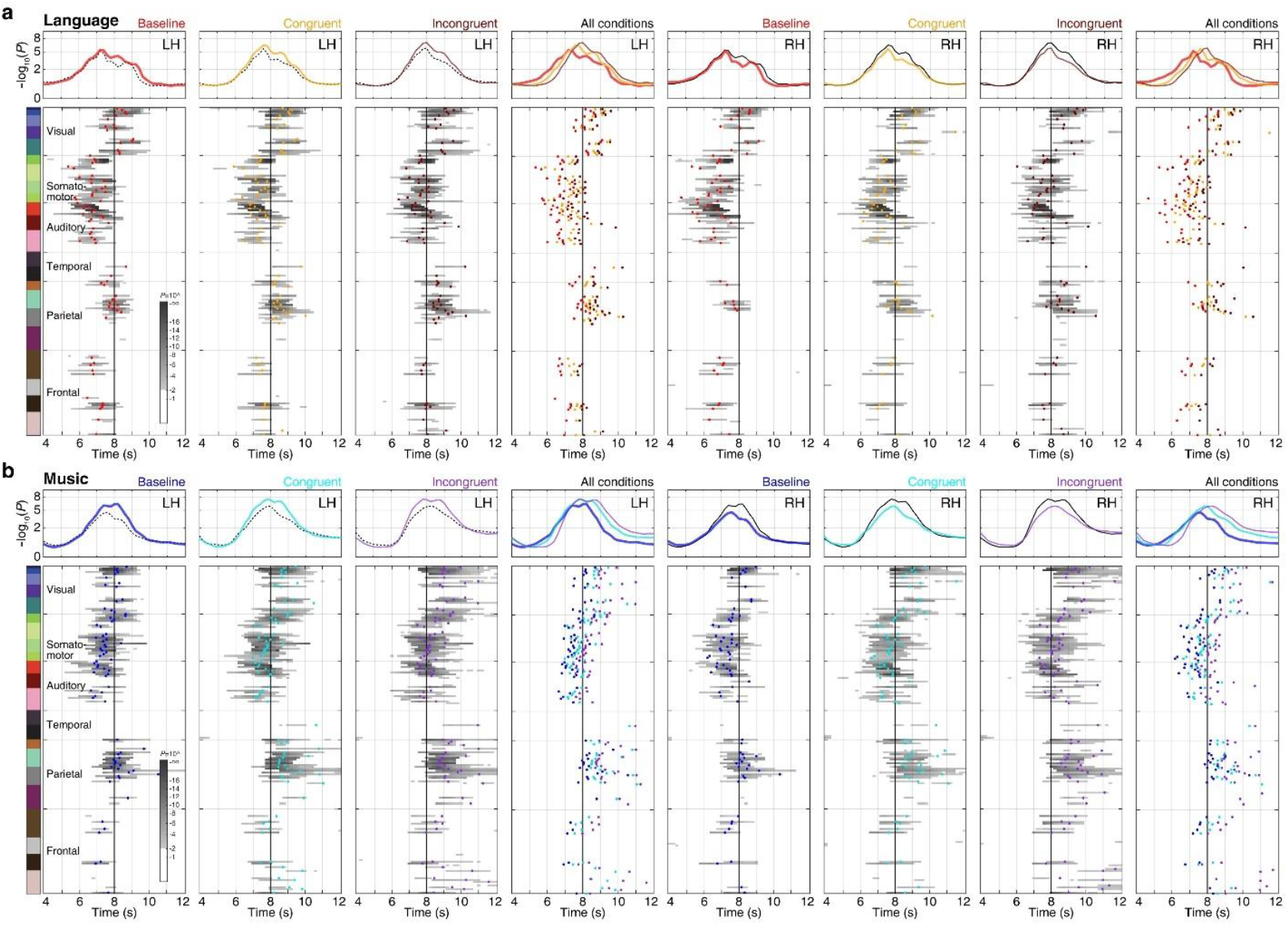
Delineating fine-grained spatiotemporal modulation of rhythm control demands. **a**, language tasks and **b**, music tasks. Surge profiles (upper panels) and Gantt charts (lower panels) illustrating activations in the left hemisphere (LH, columns 1-4) and right hemisphere (RH, columns 5-8) for baseline, congruent, and incongruent conditions in Each black (dotted or solid) curve in the surge profile panels indicates the overall phase distribution in the contralateral hemisphere for each condition. The surge height at each time point is calculated by -log_10_(*P*-value), e.g., a surge height of 2.0 indicates an SNR with *F*(2, 230) = 4.7, *P* = 0.01 (Methods). Each Gantt chart shows the surge profiles (as horizontal grayscale bars) of 180 sROIs in each hemisphere, which are displayed in ascending order according to the list of HCP-MMP1 parcellation in Supplementary Table 1. For each sROI surge profile (bar), only the portion with a surge height (SNR) above 2.0 (*P* < 0.01) is shown. The colorbar indicates 22 sections of sROIs sequentially grouped from visual to frontal cortices. The dot on each bar marks the mean phase estimated by vector-averaging across vertices within the corresponding sROI (Supplementary Table 1e). The fourth and eighth columns compare whole-hemisphere surge profiles and sROI mean phases across three conditions in each domain. In each Gantt chart, the black line at 8 s delineates earlier-phase and later-phase activations.

Relative to the baseline, congruent and incongruent conditions elicited delayed activations with higher amplitudes in the whole-hemisphere surge profiles in both domains (Fig. 5a,b, upper panels). Across all conditions, the left hemisphere consistently exhibited stronger activations than the right hemisphere in both domains. The Gantt chart further revealed moment-to-moment propagation of hemodynamic activations from one set of brain regions (sROIs) to another, as indicated by the progressive transition of the bars and sROI mean-phase markers (Fig. 5a,b, lower panels). Both language and music baseline conditions initially activated the supplementary motor, premotor, primary sensorimotor, auditory, dorsal lateral prefrontal, anterior insular, and anterior and posterior cingulate regions, followed by subsequent engagement of the visual, posterior parietal, and temporal cortices (Supplementary Movies 1–6). The same set of brain regions were consistently activated and modulated by increasing rhythm control demands in both language and music tasks, maintaining largely the same spatiotemporal formation (i.e., activation order and relative timing between sROIs) while exhibiting rightward shifts in congruent and incongruent conditions (Supplementary Movie 7). The increased activation amplitudes across conditions likewise reflected rhythmic modulation.

Hemodynamic traveling-wave movies visualize the timing, location, and direction of information flows. The evolving activity in each task can be examined by moving a vertical slider (spanning 180 sROIs) along the Gantt chart timeline (Supplementary Movies 1–6). At any given moment, the slider highlights the sROIs with significant activations, corresponding to the spatial patterns on the flattened surface in the movie frame. As the slider advances, the activation patterns — manifesting as traveling waves — transition smoothly from one set of brain regions to another. The differences in processing speed across conditions are best illustrated in the overlaid Gantt charts and surge profiles (Fig. 5a,b, fourth and fifth columns) as well as in the traveling wave movies showing spatial and temporal changes of brain activations through condition transitions in the Gantt charts (Supplementary Movie 7).

### Consistent modulations in brain activation phases and behavioral responses across domains

Fig. 6a shows the distribution of sROI mean phases in each Gantt chart in Fig. 5. Both language and music tasks show a similar tendency of temporal shifting in mean phases from baseline through congruent to incongruent conditions. The circular means of sROI mean phases in each hemisphere are significantly different between conditions within each domain (*P* < 0.05, Watson-Williams test with Bonferroni correction for three comparisons), except for the differences between congruent and incongruent language conditions in the right hemisphere.

**Fig. 6.**
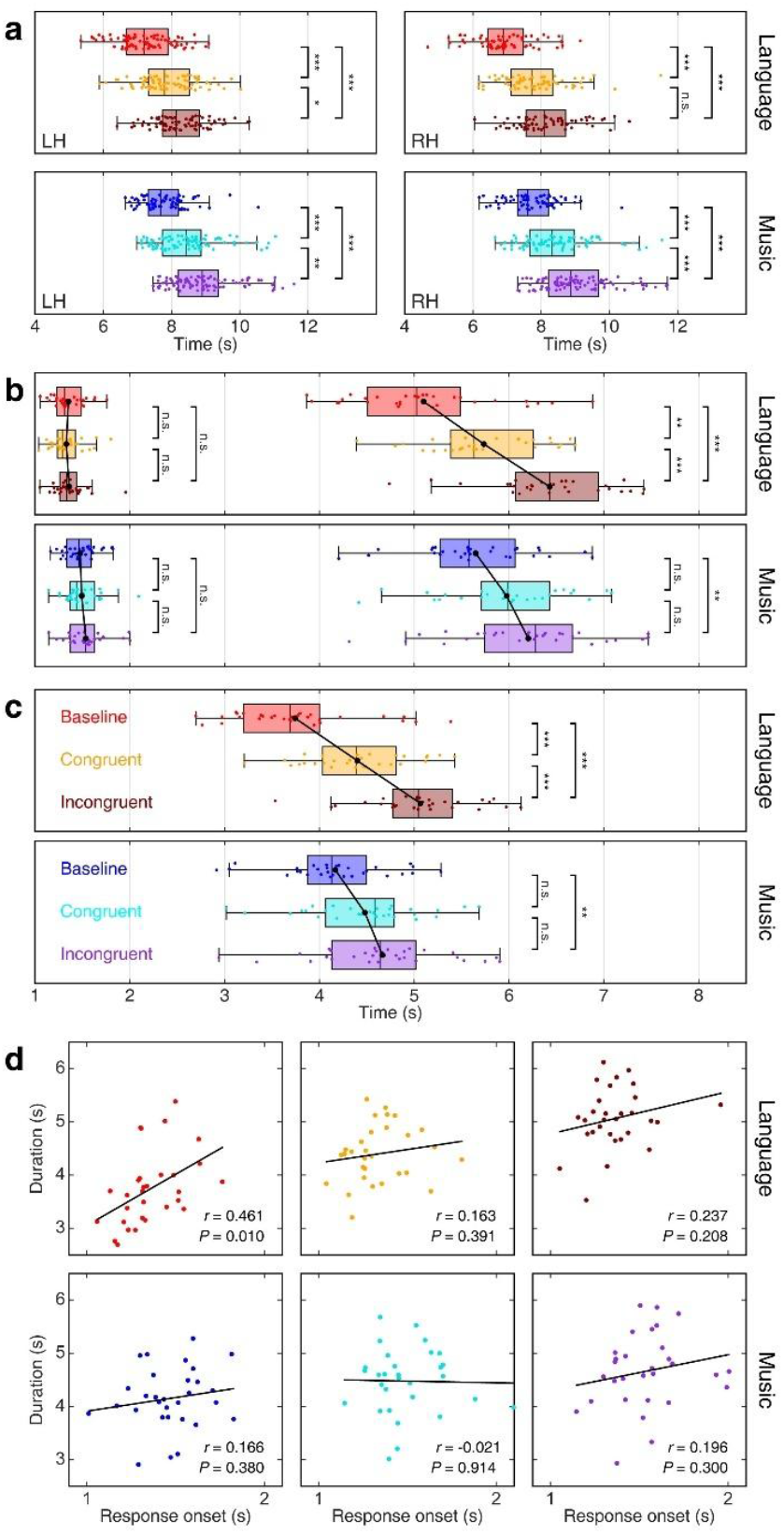
Relating task modulated activation phases to behavioral performance. **a**, Boxplots showing the group-average distribution of the mean phases across sROIs in each Gantt chart for language and music tasks (Fig. 5 and Supplementary Table 1e). LH: left hemisphere; RH: right hemisphere. **b**, Group distribution (*N* = 30) of response onsets (left boxplots) and response offsets (right boxplots) in language and music tasks. **c**, Group distribution (*N* = 30) of the response durations in language and music tasks. Color-coding schemes in all panels of Fig. 6: warm tones for language tasks (baseline: red; congruent: amber; incongruent: brown); cold tones for music tasks (baseline: blue; congruent: cyan; incongruent: purple). **d**, Scatter plots showing the correlation between response onsets and response durations in language or music tasks (*N* = 30). Levels of statistical significance in all panels: n.s., *P* > 0.05, \**P* < 0.05, \*\**P* < 0.01, \*\*\**P* < 0.001; Bonferroni corrected.

Analysis of audio recording during functional scans revealed response onsets, offsets, and durations of production in both domains (Fig. 6b,c). Response onsets in all tasks show no statistically significant differences between conditions within the language domain (*F*(2, 87) = 0.249, *P* = 0.78, *η*^2^ = 0.006; baseline: 1.36 ± 0.16 s, congruent: 1.33 ± 0.17 s, incongruent: 1.36 ± 0.17 s) and between conditions in the music domain (*F*(2, 87) = 0.756, *P* = 0.473, *η*^2^ = 0.017; one-way ANOVA, two tailed, Bonferroni, corrected for three comparisons; baseline: 1.48 ± 0.17 s, congruent: 1.5 ± 0.2 s, incongruent: 1.54 ± 0.2 s). Meanwhile, significant differences of response offsets were found in all between-condition contrasts within the language domain (*F*(2, 87) = 29.833, *P* < 0.001, *η*^2^ = 0.407; Fig. 6b). The offsets exhibited delay from the language baseline (5.1 ± 0.77 s) to congruent (5.74 ± 0.59 s), and to incongruent (6.43 ± 0.63 s) conditions. Similarly, response offsets in the music domain increased from the baseline (5.65 ± 0.61 s) to congruent (5.98 ± 0.64 s), and to incongruent (6.21 ± 0.76 s) conditions (*F*(2, 87) = 5.107, *P* = 0.008, *η*^2^ = 0.105). However, only the difference between the music baseline and incongruent conditions was statistically significant (*P* < 0.01).

Statistically significant differences in response durations were found between all contrasts of language conditions (*F*(2, 87) = 36.68, *P* < 0.001, *η*^2^ = 0.458, one-way ANOVA, two tailed, Bonferroni, corrected for three comparisons; Fig. 6c). Response durations increased from the language baseline condition (3.75 ± 0.68 s) to congruent condition (4.4 ± 0.54 s), and to incongruent condition (5.07 ± 0.57 s). The music domain showed a similar pattern in response durations that increased from the baseline (4.17 ± 0.57 s) to congruent (4.48 ± 0.62 s), and to incongruent (4.67 ± 0.7 s) conditions (*F*(2, 87) = 4.78, *P* = 0.011, *η*^2^ = 0.1). Significant difference was only found between music baseline condition and incongruent condition in pairwise comparison (*P* < 0.05).

A significant correlation between response onsets and durations was specific to the language baseline condition (Pearson’s *r* = 0.461, 95% confidence interval = [0.121, 0.704], *P* = 0.01, *n* = 30) (Fig. 6d). The absence of this correlation in other tasks indicates that the timing of response initiation does not directly predict production completion time, especially when explicit control over rhythm is involved.

Overall, the mean phases in brain activations and behavioral responses exhibited a consistent trend of delays across conditions in the language and music domains, corresponding to increasing rhythm control demands. Furthermore, significant differences in brain activation mean phases were observed between conditions in both domains. In the music domain, however, response durations showed no significant difference between the baseline and congruent conditions or between the congruent and incongruent conditions. This pattern indicates subtle distinctions in how rhythmic complexity modulates brain activation phases and behavioral response times between language and music, with language processing showing a closer alignment between brain activity and behavioral output.

### Increasing rhythm control demands induced temporal shifts in behavioral and hemodynamic responses

In a representative subject, we demonstrate that the temporal shifts in response onsets, offsets, and durations across conditions are consistent with the group behavioral patterns (Fig. 7a,b). Response onsets show no significant difference between conditions in both domains (*P* > 0.05, one-way ANOVA, two-tailed, Bonferroni, corrected for three comparisons). In contrast, nearly all response offsets and durations differ significantly between conditions in both domains (*P* < 0.05), except those between the music baseline and congruent conditions.

**Fig. 7.**
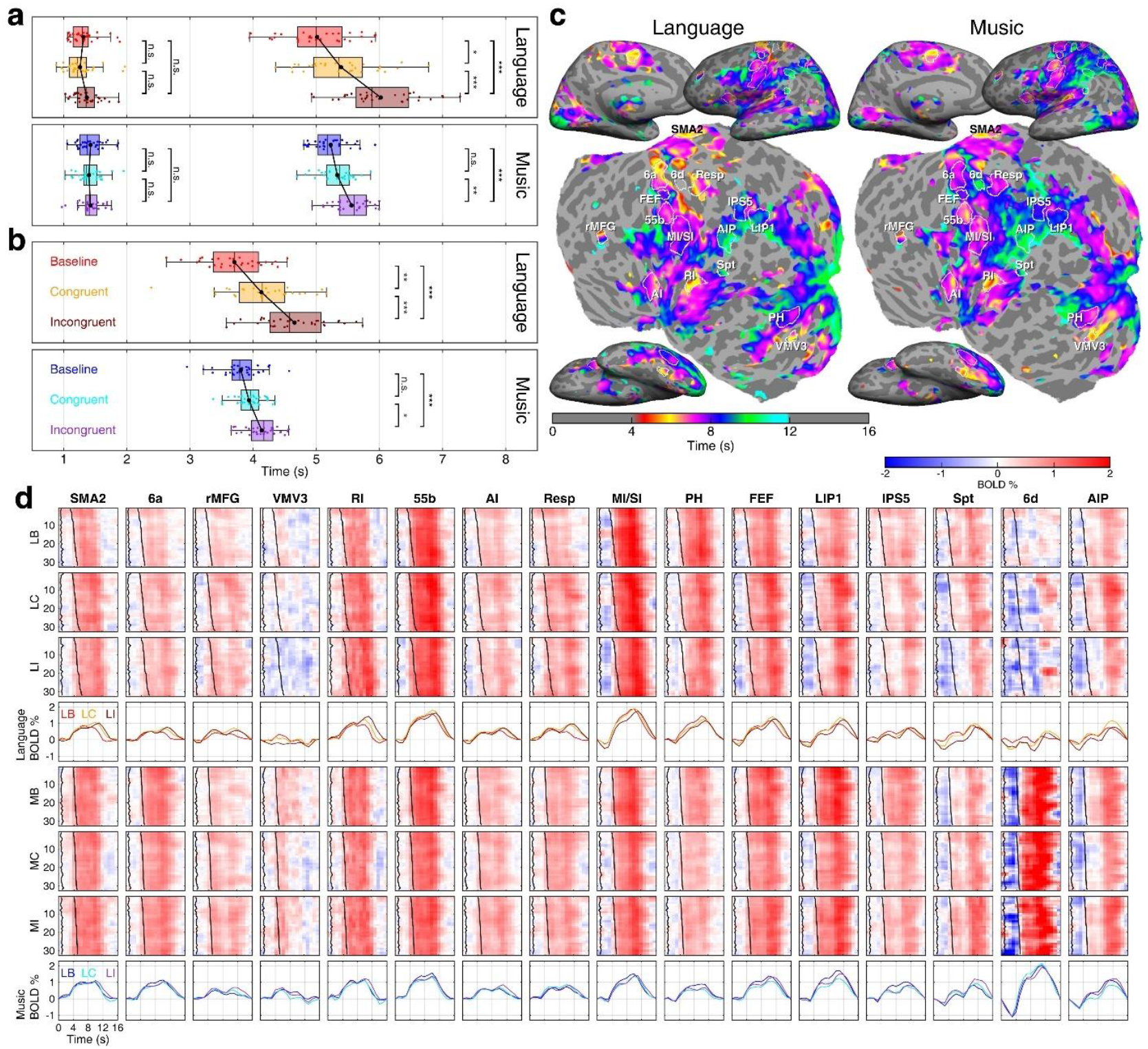
Modulations in behavioral performance and hemodynamic responses in a representative subject. **a**, Boxplots showing the distribution of response onsets (left boxplots) and response offsets (right boxplots) in language and music tasks. **b**, Boxplots showing the distribution of response durations in language and music tasks. Color-coding schemes for tasks follow those of Fig. 6. **c**, Maps of phase-encoded activations for baseline language and music tasks, overlaid with contours of selected sROIs. Levels of statistical significance: n.s., *P* > 0.05, \**P* < 0.05, \*\**P* < 0.01, \*\*\**P* < 0.001; Bonferroni corrected. **d**, Two-dimensional visualization for hemodynamic responses (BOLD images) and average time courses (BOLD curves) of 32 trials in each of 16 sROIs for each language or music task. In each BOLD image, the first black curve indicates response onset, and the second indicates response offset. Each BOLD curve was obtained by averaging the hemodynamic responses across 32 trials in a 16-s cycle for each condition. LB, LC, and LI: baseline, congruent, and incongruent conditions in language tasks; MB, MC, and MI: baseline, congruent, and incongruent conditions in music tasks.

To investigate how language or music production dynamically engages the brain under different rhythm control demands, we analyzed trial-by-trial hemodynamic responses in 16 sROIs selected from several brain regions of the representative subject (Fig. 7c). These sROIs include motor planning regions (SMA2, 6a, and rMFG), ventral visual areas (VMV3 and PH), auditory regions (AI, RI, and 55b), orofacial sensorimotor cortex (MI/SI), respiratory control area (Resp), eye movement control areas (FEF, LIP1, and IPS5), manual control areas (6d and AIP), and an auditory-motor interaction area (Spt).

The blood-oxygenation-level-dependent (BOLD) signals of 32 trials of each condition are sorted by response offsets and displayed as a BOLD image for each sROI. The BOLD image in each condition exhibits a gradually shifting trend that matches the sorted response-offset curve. The average BOLD curves in each sROI show systematic shifting patterns across conditions. Each sROI exhibits a unique BOLD profile, with some engaged earlier and others later, suggesting that they engage like different sections of an orchestra during language or music production. Specifically, SMA2, 6a, 55b, RI, AI, Resp, and MI/SI engage earlier and are primarily involved in the preparatory functions of rhythm planning. Other sROIs, including PH, FEF, LIP1, IPS5, Spt, 6d, AIP, and rMFG, are engaged later and contribute to the execution and monitoring of production. In addition, area 6d shows strong hemodynamic responses in music but not language tasks, reflecting its role in manual control during music production.

Across domains, the overall BOLD curves show consistent temporal shifts in their onsets, peaks, and offsets, indicating a clear pattern of rhythm-dependent modulations (Fig. 7d, rows 4 and 8). Together, these spatiotemporal activation patterns reveal how different cortical regions coordinate as ‘neural ensembles’ to support complex behavioral performance in both domains. Each region plays a distinct yet dynamically coordinating role, collectively orchestrating language or music production under varying rhythmic contexts, giving rise to behaviors with the underlying neural dynamics that unfold in harmony.

## Discussion

Direct comparison between the neural mechanisms underlying overt language and music production has been challenging for noninvasive neuroimaging, particularly fMRI. Overcoming the technical limitations imposed by head motion and scanner noise, we used rapid phase-encoded fMRI to capture brain activations with both high spatial and high temporal resolutions during speaking and wind instrument playing. The spatiotemporal brain dynamics revealed in the current study have not been accessible through continuous imaging of covert production tasks^42^ and sparse sampling of intermittent activations in overt production tasks ^16,22^. Furthermore, while continuous brain activity can be recorded during overt speech production, previous noninvasive neuroimaging studies have predominantly used tasks involving picture naming or single-vowel (or word) utterances^43-47^. In the current study, rapid phase-encoded fMRI added an extra dimension, time, to static statistical maps, revealing where and when hemodynamic traveling waves propagate across the cortical surface during the continuous overt production of language and music sequences.

Apart from the technical challenges, comparing language and music processing on common ground necessitates careful alignment of tasks and stimuli between domains. The current study employed naturalistic and real-time production tasks involving reading aloud sentences and recorder playing of musical phrases, which are thought to be the most comparable forms of language and instrumental performance. Unlike common bimanual instruments such as the piano, recorder playing involves orofacial, vocal tract, and respiratory control during the production of sequences with various rhythmic patterns, making recorder playing more directly comparable to speaking. At the same time, it remains distinct from singing, which closely resembles speech production rather than functioning as an instrumental form.

By analyzing surface traveling wave patterns in phase-encoded fMRI data, we showed that speaking and recorder playing engaged largely shared but partially distinct spatiotemporal neural streams. While the activation timing consistently shifted across conditions in both domains, the shared streams were differentially tuned by rhythmic complexity, with music tasks engaging additional language areas along with increased rhythm control demands. Specifically, tasks in both domains induce overlapping neural flows that unfold over time in a similar formation of traveling waves, transitioning from one set of brain regions to another (Fig. 5 and Supplementary Movies 1-6). These dynamic spatiotemporal patterns highlight how hemodynamic activity progresses across the brain, which is unobtainable using conventional contrast-based fMRI designs and volume-based analyses. Crucially, the overlapping streams are modulated by rhythm in both spatial and temporal dimensions. As rhythm control demands increase, shared regions not only grow spatially, with music related activations expanding into language areas in congruent and incongruent conditions, but also shift temporally, with activations occurring later in these conditions. These rhythm-sensitive modulations reveal that the interaction between music and language processing is not fixed, but dynamically shaped by task complexity, which becomes most evident when the full spatiotemporal flow patterns are considered across the baseline, congruent, and incongruent conditions (Figs. 5, 6; Supplementary Movie 7).

Previous studies have found neural overlaps between language and music processing in the temporal lobe, including the Heschl’s gyrus, planum temporale, planum polare, STS, and STG^48-50^, which are responsible for the acoustic analysis of speech and music involving phonological decoding and prosody processing^51,52^. Extending these findings, we observed stronger activations in the middle to posterior STS and area Spt in the left hemisphere (overlapping areas PF, PFcm, and PSL in the HCP-MMP1 atlas) during speaking than during recorder playing, suggesting language-specific functions in these regions^53,54^. Furthermore, in the current study, extensive overlaps between language and music processing were observed in the premotor cortex as well as orofacial and respiratory representations in the sensorimotor cortex, which support motor planning and execution during the oral production of linguistic and musical sequences^48,49^. Additionally, recorder playing elicited bilateral activations in manual control regions in the premotor, sensorimotor, and posterior parietal cortices. In the frontal cortex, the middle and inferior frontal gyri were commonly activated in both domains, reflecting shared mechanisms for hierarchical structure processing and higher-order cognitive control^34,55^. Compared with music tasks, however, language tasks recruited more extensive areas within IFG, suggesting domain-specific demands in linguistic processing.

Turning to rhythm processing, the overlapping regions activated by language and music in this study align with the fronto-parietal, SMA–parietal, and auditory–motor networks, which have been suggested to support rhythm retrieval, temporal movement preparation, and rhythm prediction across both domains^8,13-35,56,57^. Moreover, previous studies have linked these regions to the processing of rhythmic patterns that vary in complexity and irregularity^18,22,27^. While prior research has primarily focused on static cortical overlaps between language and music processing, the dynamic interaction among these regions has remained a key piece missing from the whole picture of rhythm production. Using a rapid phase-encoded design, the current study continuously captured simultaneous and sequential activations among brain regions during naturalistic rhythm control tasks in both domains. In the conjunction maps between domains, we identified expansion of neural overlaps in conditions with conscious rhythm control, during which music recruited more language-dominant areas, including IFG, STS, and insula (Figs. 3, 4). This aligns with previous findings and suggests the recruitment of language ‘instruments’ during music production with rhythms of different complexities. For example, in the frontal lobe, the processing of metrical structures in music largely recruited a language area, BA47, indicating that it is less domain-specific than previously assumed^29,30^. Furthermore, the Broca’s area is known to be involved in language syntactic processing with deviations from regular rhythmic patterns^21^. Similarly, the expanded overlap observed in the insula suggests stronger interactions between language and musical rhythm, reflecting shared mechanisms of temporal hierarchy processing and predictive coding across the two domains^20,22^. In the middle temporal lobe, we identified expanded overlaps in TOPJ1 and PHT under rhythm control conditions compared to the baseline music task. This expansion was associated with increased activations in congruent and incongruent music conditions, suggesting that incorporating rhythm engages the temporal regions more strongly. This pattern aligns with prior evidence that the primary auditory cortex, planum temporale, and superior temporal regions are sensitive to temporal variation in both language and music^58-60^. Overall, the expansion of activations into language-dominant regions in music tasks suggested that the spatiotemporal dynamic patterns changed along with increased rhythm control demands within the shared neural resources between language and music production (Fig. 3). The expanded overlapping areas are closely associated with the domain-general system for temporal parsing, structure prediction, and error monitoring, which are essential components in complex language and music processing. Conscious control over producing structural sequences requires increased shared demands in resolving expectation violations and attention allocation in both domains.

Beyond shared neural resources, we further identified common modulations of rhythm processing related to different metrical demands in the language and music domains. Behavioral and neural patterns across language and music tasks followed a consistent trend (Fig. 6): relative to the baseline condition, congruent conditions showed intermediate response times, durations, and activation phases and amplitudes, while incongruent conditions showed the slowest and longest behavioral responses as well as the most delayed and pronounced brain activations, as indicated in hemodynamic surge profiles (Fig. 5). Based on both behavioral and neuroimaging data, we infer that higher rhythm predictability and regularity facilitate sequence processing. In the congruent condition, where explicit regular accent symbols are present, the brain is sufficiently activated to support rapid and prediction-based production. This condition is more engaging than the baseline condition, which involved minimal conscious rhythm processing and produced lower activations. At the same time, the congruent condition is less demanding than the incongruent condition, which required greater effort and resulted in slower response times, extended response durations, and delayed activations. These findings align with previous research suggesting that rhythmic predictability enhances processing efficiency^61-64^. In contrast, rhythm irregularity imposed higher demands on working memory, focused attention, and self-monitoring, thereby increasing the recruitment of neural resources and lengthier processing time. For example, previous studies have indicated that increased rhythmic complexity is associated with greater activation in the right IFG and STG, highlighting additional demands on auditory-motor integration^17,18^. Similar activation patterns have been observed in simple singing tasks, where engagement of the bilateral pars orbitalis (BA47), insula, and left cingulate gyrus was associated with rhythmic complexity^22^. Likewise, in finger-tapping tasks, pre-SMA and SMA, PMC, and DLPFC activities were proved to be correlated with rhythmic complexity^27^.

Our findings extend previous research by showing that rhythmic complexity modulates information flows through shared language–music networks, suggesting that the two domains rely on overlapping pathways but operate at different timings depending on processing demands. Moreover, while both brain activity and behavioral production in language and music tasks follow a systematic time delay across conditions, behavioral responses in the music domain show less significant differences between conditions than those in the language domain (Fig. 6). These results suggest subtle differences in the modulations between the language and music domains, as well as between neural and behavioral responses, which should be investigated through targeted experimental studies in the future.

With the spatiotemporal activation patterns revealed by rapid phase-encoded fMRI, this study provides empirical evidence for the link between language and music production, complementing existing theories and models that have conceptually outlined the connection between auditory and motor systems based on anatomical projections^8,9^. In the neural ensemble maps (Figs. 2, 3), regions exhibiting the same activation phase suggest that they coordinate in harmony and are functionally connected with each other. The moment-to-moment propagation patterns in the Gantt chart and traveling wave movies further reveal the direction of neural information flows, from one set of regions to another (Fig. 5 and Supplementary Movies 1-6). Regions with earlier phases are likely engaged in auditory-motor coupling that enables real-time anticipation and planning of upcoming events, and regions with later phases reflect domain-specific production in both domains. These findings align with the ASAP^8^ hypothesis, which posits that the motor system internally simulates auditory input to support precise temporal prediction. In line with PRISM^9^, our findings suggest that both language and music rely on the recruitment of motor planning regions to anticipate the timing and structure of incoming sequences.

The observed spatiotemporal activations can also be interpreted with dual-stream models^10,11^, which emphasize the transformation of sensory input into motor output. Beyond auditory–motor coupling, we identified the visual and posterior parietal cortices as well as Spt^38,42^, which exhibited later activation phases in both domains. Located in the dorsal auditory stream of the dual-stream models, Spt is part of the pathway connecting posterior temporal regions with premotor and parietal cortices, functioning as a hub for online monitoring of auditory-motor coupling, and supporting motor planning and execution in both language and music production. Moreover, area Spt is located at the posterior lateral sulcus (Sylvian fissure), a region also found to respond to visual (motion), auditory, and somatosensory stimulation in topological mapping studies^65,66^. The later activation phases found in the visual cortex, posterior parietal cortex, and Spt suggested crossmodal coordination during real-time reading aloud (language) and sight-reading (music) tasks.

Overall, this study illustrates dynamic response patterns in neural dynamics from predictive simulation to output execution for language and music rhythm production. Both language and music production engage a coordinated interplay among distributed brain regions, as reflected by the distinct temporal profiles featuring the rise, peak, and fall of BOLD signals across sROIs (Fig. 7) — functioning, in a sense, like different sections of an orchestra. Moreover, the BOLD profiles are systematically modulated by rhythm control demands, highlighting how rhythmic complexity shapes the recruitment and timing of shared neural resources in both domains. As different brain regions play their parts, they collectively orchestrate the precise timing needed for language and music production.

Crucially, under conscious rhythm control in congruent and incongruent conditions, the overlap between language and music processing networks became more pronounced. This suggests that both domains draw on a common architecture for temporal sequencing and prediction, and that rhythm control demands push these systems toward optimal coupling. The shift from less to greater overlaps along the streams across conditions implies a dynamic allocation of neural resources depending on cognitive load and coordination demands. Under conditions of explicit rhythm control, the network of language and music regions operates in higher synchrony, akin to an ensemble performing in unison.

A limitation of this study is that participants’ brain responses in the language tasks might have been partially influenced by their prior knowledge of English pronunciation. However, this very familiarity also allowed the tasks to effectively distinguish between expected accent positions in the congruent condition and unexpected ones in the incongruent condition, thereby paralleling musical predictability in terms of the hierarchical organization of accent positions in the brain. Alternative approaches, such as using pseudowords or employing a language completely unknown to participants, would have made the production of sequences more akin to singing than to speaking, as participants would be articulating random units without linguistic meaning. For the same reason, such approaches would not provide a fair parallel to the musical stimuli either, given that participants already possess structured knowledge of musical notes. Future studies could further disentangle the influence of linguistic familiarity and metrical predictability by systematically manipulating the degree of stress regularity within a familiar language, and by comparing the brain activation and behavioral performance of native speakers, non-native speakers, and participants entirely unfamiliar with the same stimuli. Such designs would help clarify whether the observed neural effects primarily reflect sensitivity to rhythmic predictability shaped by prior linguistic experience.

## Conclusion

By investigating overt language and music production, two uniquely human capabilities, through parallel tasks and phase-encoded fMRI, this study enables a direct within-subject comparison of the spatiotemporal brain dynamics during speaking and wind instrument performance. The findings reveal both common and distinct neural mechanisms supporting language and music production. Specifically, speaking and recorder playing show overlapping activity in auditory–motor regions during earlier phases, while later phases involve shared engagement of visual and posterior parietal cortices and area Spt, suggesting a common neural ensemble supporting crossmodal integration, structural prediction, and error monitoring. For both domains, increasing rhythmic complexity modulated activation timing and amplitude, with more predictable rhythms eliciting faster but weaker responses, and irregular rhythms producing slower, stronger activations. Explicit rhythm control in wind playing recruited additional ‘neural instruments’ typically engaged in language processing, indicating that rhythmic predictability and motor output jointly shape the neural coordination underlying rhythm control. Together, these results deepen our understanding of the neural correlates of structured sequence production in language and music and open new avenues for investigating how the brain flexibly integrates linguistic and musical actions under varying rhythmic demands.

## Methods

### Participants

Thirty native Cantonese speakers (22.1 ± 5.92 years old; 21 females, 9 males) who were amateur wind instrument players participated in this study. All subjects spoke English as their second language (self-reported average proficiency score = 0.62 ± 0.03 on the Language History Questionnaire^67^). Participants had 12.07 ± 4.07 years of formal music training. They were all right-handed, had normal to corrected-to-normal vision, and had no history of neurological impairment.

### Experimental design

In each fMRI session, the subject read stimuli presented on a screen and performed three language tasks and three music tasks, with baseline, congruent, and incongruent conditions in each domain (Fig. 1c). Each task was repeated twice in nonconsecutive 256-s scans, each of which comprised sixteen periodic cycles. Within each 16-s cycle, the subject spoke a sentence or played a musical phrase during the task period (0-6 s) and viewed a blank screen during the rest period (6-16 s).

The language stimuli included 32 English sentences, each consisting of 13-14 syllables. The procedures for creating language stimuli followed those of a previous study^41^, ensuring that the sentences were natural and authentic by drawing examples from Collins Dictionary English (https://www.collinsdictionary.com/dictionary/english/corpus). Moreover, all words used in the sentences were part of *Longman Communication 3000*, a collection of the 3,000 most frequently occurring words in spoken and written English, which ensured readability for participants. Each sentence inherently contained five naturally occurring accent positions. The music stimuli included 32 phrases of diatonic tones presented in staves. Each musical phrase consisted of 10 eighth notes in C major, randomized between C3 and B3.

In each language or music task, the subject spoke sentences or played musical phrases under one of three conditions of rhythm control demands, where stimuli were presented: (1) without accent symbol (baseline condition), (2) with regularly placed accent symbols (congruent condition; accent symbols placed on natural and regular accented positions in sentences and on downbeat positions in musical phrases), or (3) with irregularly placed accent symbols (incongruent condition; two to three out of the five accents placed on unnatural or irregular positions in sentences and on downbeat positions in musical phrases). All subjects were familiar with the classical accent symbol “>” in Western musical notation.

### Experimental setup

Every subject underwent a training session to keep his or her head still while speaking and playing a recorder in an MRI simulator (Shenzhen Sinorad Medical Electronics Inc., China) prior to an fMRI session. Head motion was monitored in real time by a motion sensor attached to the forehead (MoTrak, Psychology Software Tools, Inc., Sharpsburg, PA), which triggered alarms when movement exceeded thresholds (>1 mm displacement or > 1º rotation). For head immobilization during overt production tasks in the MRI scanner, subjects wore a custom-fitted facial mask made of a thermoplastic sheet (Fig. 1a; Sun Medical Products Co., Ltd., China). Auditory feedback was provided via a pair of MR-compatible headphones during speaking and recorder playing in the MRI simulator. During the actual fMRI experiment, subjects lay supine within a head coil filled with deformable resin clay and wore earplugs along with MR-compatible noise-cancellation headphones (OptoActive II, OptoAcoustics Ltd., Israel) (Fig. 1b). A rear-view mirror mounted on the head coil enabled visualization of visual stimuli presented on a 40-inch MR-compatible LCD monitor (InroomViewingDevice, NordicNeuroLab AS, Norway) located behind the scanner. A soprano recorder (German TRS-23, Yamaha Corp., Japan) was positioned above the participant’s chest. An extended mouthpiece was attached to the recorder to allow proper positioning and manual control outside the head coil (Fig. 1b). The extension was constructed using plastic modular hoses (Loc-Line Inc., Lake Oswego, OR), which permitted flexible adjustment of its angle and length. Experiment Builder (SR Research Ltd.) was used to present stimuli and record event timings. Each functional scan was initiated by a synchronization signal (“s” key) from the SyncBox (NordicNeuroLab AS). Speech output and recorder music were captured using an MR-compatible microphone (OptoAcoustics Ltd.) and recorded continuously during each functional scan using OptiMRI 3.1 Software (OptoAcoustics Ltd.).

### Behavioral data analysis

The timings of voice or recorder output were manually annotated from the audio recordings using Audacity software (https://www.audacityteam.org). To quantitatively characterize the temporal dynamics of behavioral responses under the baseline, congruent, and incongruent conditions, we extracted three key temporal metrics for each trial: onset time (latency between stimulus onset and response initiation), offset time (latency between stimulus onset and response completion), and response duration (the difference between onset and offset times).

For each subject, the parameters were averaged across 32 trials in two scans of the same condition, and were further averaged across subjects for each condition. A one-way repeated-measures analysis of variance (ANOVA) was conducted to examine the effect of the three conditions on behavioral responses in each domain, followed by Bonferroni-corrected post hoc pairwise comparisons. Furthermore, we computed Pearson correlation between onset time and response duration across 30 subjects (Fig. 6d).

### Image acquisition

All brain images were scanned with a 32-channel head coil in a Siemens MAGNETOM Prisma 3T scanner at the Centre for Cognitive and Brain Sciences, University of Macau. Each fMRI session included twelve functional scans and two structural scans. For each subject, twelve scans of functional images were acquired using a single-shot echo planar imaging (EPI) sequence with blipped-CAIPIRINHA simultaneous multi-slice (SMS) reconstruction. The parameters for the EPI sequence were: acceleration factor = 5, TR = 1000 ms, TE = 30 ms, flip angle = 60º, 55 interleaved ascending axial slices, field of view = 192×192 mm, matrix = 64×64, voxel size = 3×3×3 mm, bandwidth = 2368 Hz/Px, 6 dummy TRs, and 256 TRs per scan. Two sets of structural images were acquired using an MPRAGE sequence with the following parameters: TR = 2300 ms, TE = 2.26 ms, TI = 900 ms, flip angle = 8º, field of view = 256×256 mm, slice thickness = 1 mm, 256 axial slices, matrix size = 256×256×256, voxel size = 1×1×1 mm, bandwidth = 200 Hz/Px, 234 s per scan. Functional and structural images were prescribed with identical slice center and orientation for direct alignment between them.

### Image preprocessing

All raw (.ima) images were first converted to Analysis of Functional NeuroImages (AFNI; https://afni.nimh.nih.gov/) BRIK files by the AFNI *to3d* command. Functional BRIK files were motion-corrected and registered with the first volume of the seventh functional scan (the middle volume among twelve scans) using the AFNI *3dvolreg* command. The motion-corrected functional images then underwent slice-timing correction and coregistration with structural images via the *csurf* package^66^ (https://pages.ucsd.edu/~msereno/csurf/). For each individual subject, two sets of structural images were averaged and then subjected to cortical surface reconstruction in FreeSurfer 7.2 (https://surfer.nmr.mgh.harvard.edu/)^70,71^.

### Functional data analysis

Following motion and slice-timing correction, functional images underwent a data processing pipeline consisting of voxel-wise Fourier analyses and vertex-wise (surface-based) complex analyses developed by Sereno and colleagues^66,68-75^. These analyses are standard functions included in the *csurf* package. For each functional scan (64×64×55×256 data points), the time series *x*_*m*_(*t*) of each voxel *m* was analyzed with a 256-point discrete Fourier transform:

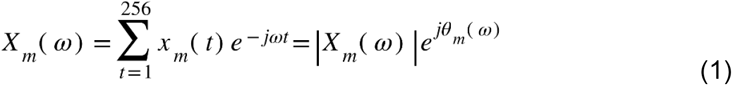

where *X*_*m*_*(ω)* is the Fourier component at frequency *ω* between 0-127 cycles per scan, and |*X*_*m*_*(ω)*| and *θ*_*m*_*(ω)* are its amplitude and phase. The task frequency is defined as *ω*_*s*_ (16 cycles per scan), at which the BOLD signal fluctuates periodically in response to periodic stimuli and tasks. The signal 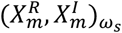 and noise 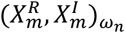 in voxel *m* are the Fourier components at task frequency *ω*_*s*_ and other non-task frequencies at *ω*_*n*_, respectively. The statistical significance of the signal-to-noise ratio (SNR)^40,41,68,69,74-77^ in voxel *m* was evaluated by:

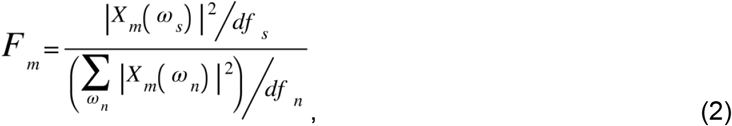

where *df*_*s*_ = 2 (degrees of freedom of real and imaginary components at *ω*_*s*_ = 16 cycles per scan) and *df*_*n*_ = 230 (degrees of freedom of real and imaginary components at non-task frequencies *ω*_*n*_: excluding 0-3, 15, 17, 31-33, 47-49, and 64 from [0, 127] cycles per scan, i.e., *df*_*n*_ = (128 -4 -2 -3 -3 -1)*2 = 230). The *P*-value of the *F*-ratio in equation (2) was estimated by the cumulative distribution function *F*(*F*_*m*_; *df*_*s*_, *df*_*n*_)^40,41,75-77^. A complex *F*-value,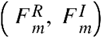, incorporating both the *F*-value and phase, *θ*_*m*_*(ω*_*s*_), of the periodic signal in each voxel was obtained by:

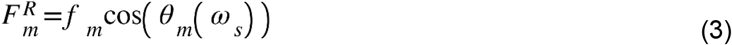

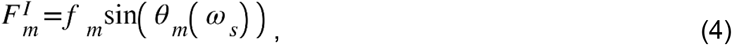

where *f*_*m*_ is the square root of *F*_*m*_.

For each subject *S*, the complex *F*-values in the corresponding voxel *m* were vector-averaged (voxel-wise) across two scans, k ={1, 2}, for the same language or music task by:

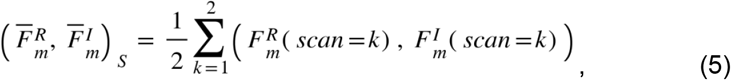

using the “Combine 3D Phase Statistics” function in *csurf*. The resulting subject-level voxel-wise average *F*-values, 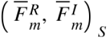, were then projected onto vertex *v* on the individualized cortical surfaces of subject *S*, yielding an average 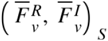 map for each task.

The spherical averaging method^72-77^ (“Cross Session Spherical Average” function in *csurf*) was used to obtain surface-based group-average maps for each task. First, each average map of a single subject, 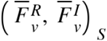, was resampled to a common spherical coordinate system using the FreeSurfer *mri_surf2surf* command (https://freesurfer.net/fswiki/mri_surf2surf). Second, the complex *F*-values of each vertex *v* on the common spherical surface were vector-averaged (vertex-wise) across subjects (*N* = 30) by:

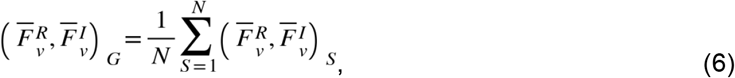

which yielded a map of group-average complex values, 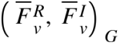, for each task. The amplitude and phase of each vertex in the group-average maps were obtained by:

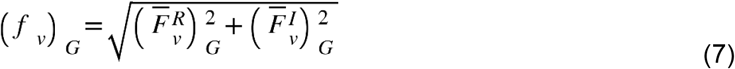

and

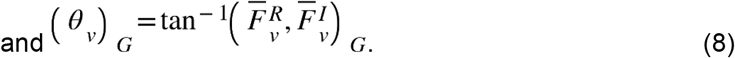

The amplitudes at each vertex in the common spherical coordinate system were tested across subjects by *F*-statistics (*n* = 30, *F*(2, 40) > 5.18, *P* < 0.01) and corrected for multiple comparisons using a surface-based cluster-size exclusion method^73^; cluster size = 64 mm^2^, alpha = 0.05, corrected). Finally, the amplitude and phase of each vertex in the cluster-corrected group-average maps were displayed on the inflated and flattened cortical surfaces of *fsaverage* using the *csurf* package (Figs. 2, 3). The activation phase (time delay) at each vertex *v* was color-coded according to a colorbar representing a continuous progression of phases, containing task-positive activations.

### Conjunction analysis

To compare the spatial extent of activations between domains, we searched for vertices with statistically significant activations (n = 30, *F*(2, 58) = 4.99, *P* < 0.01, cluster corrected) that were specific to the language task, specific to the music task, or common to both under each condition. The results are displayed as surface-based conjunction maps on the flattened cortical surfaces of *fsaverage* (Fig. 3), with different colors indicating task-specific and shared activations. The percentages of vertices activated by a single task or both tasks are summarized in Table 1.

### ROI analysis

To assess the differences in activation patterns across tasks, we used the HCP-MMP1 atlas^78^ to parcellate the *fsaverage* cortical surface into 180 surface-based regions of interest (sROIs) in each hemisphere (Supplementary Fig. 1a). For each sROI, we quantified the ratio between the significantly activated vertices (Fig. 1b, *P* < 0.01, cluster corrected) and the total number of vertices within it, yielding *Q*_LH_ and *Q*_RH_ for each pair of corresponding sROIs in the left and right hemispheres, respectively. We then computed a laterality index (LI)^79^ for each condition to measure hemispheric asymmetry by:

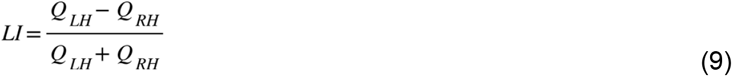

The results are shown in Supplementary Fig. 1c and Supplementary Table 1d. For each task in the language and music domains, we further computed the ratios of vertices with earlier-phase (4-8 s) and later-phase (8-12 s) activations in each sROI (Fig. 4b, Supplementary Figs. 2-4, and Supplementary Table 1b).

### Hemodynamic surge profiles and Gantt charts

For each task, we analyzed the distribution of activation phases of 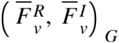 of all vertices within each hemisphere and each sROI. The results are presented as hemodynamic surge profiles^41^ and Gantt charts (Fig. 5). First, the complex plane was divided into 81 equally spaced bins between 0° (0 s) and 360° (16 s) at *d* ={0°, 4.5°, 9°, …, 360°}, equivalent to {0, 0.2, 0.4, …, 16 s}. Second, for each hemisphere or each sROI, a total of *D* vertices were identified with their phases, *(θ*_*v*_)_*G*_, falling within a moving sector (range = 9° [0.4 s]; step = 4.5° [0.2 s]) centered at each bin *d*. Third, the complex *F*-values of *D* vertices within the moving sector [*d*-4.5°, *d*+4.5°] were averaged by:

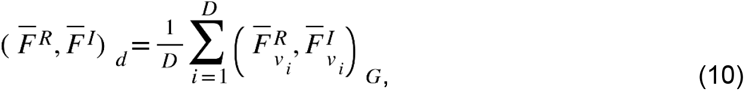

and the average magnitude was obtained by:

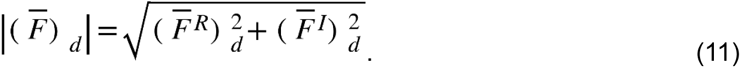

Lastly, a *P*-value, was estimated using the cumulative distribution function, 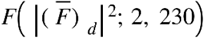, which represents the average SNR of hemodynamic surge at each time *d*. The height of a surge profile^41^ at each time point *d* is obtained by -log10(*P*), as shown in the y-axis of the upper panels of Fig. 5a,b.

A Gantt chart was created for each task by displaying the surge profiles of 180 sROIs in each hemisphere as grayscale bars (Fig. 5a,b, bottom panels). Each grayscale bar represents the time interval during which the surge height exceeds 2.0, corresponding to an SNR with *P* < 0.01. Portions with surge heights below 2.0 are colored white, indicating either subthreshold or absent responses. All surge profiles and Gantt charts are shown for a time window between 4 and 12 s, encompassing the full span of significant hemodynamic responses associated with the tasks. Each dot marker on a grayscale bar indicates a surge profile’s mean phase, *θ*_*sROI*_, obtained by averaging the complex *F*-values of *V* vertices within each sROI by:

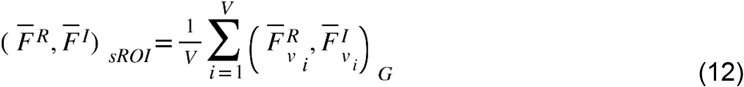

and

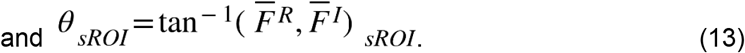

### Circular statistics

Fig. 6a presents the distribution of mean phases (*θ*_*sROI*_) across all sROIs in each hemisphere for each task. We applied the Watson Williams test for circular data^80-82^ to examine whether the circular means of sROI mean phases differed significantly between tasks. Given the mean complex *F*-values of *n*_*1*_ sROIs for Task 1 and *n*_*2*_ sROIs for Task 2:

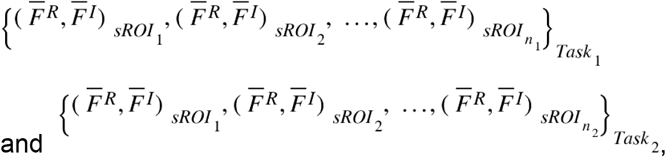

and let *n* = *n*_*1*_+*n*_*2*_, an *F*-value of the Watson-Williams test is obtained by:

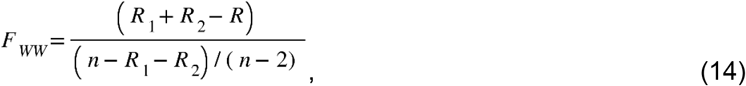

where *R*_*1*_, *R*_*2*_, and *R* are computed from the radian representations^81^:

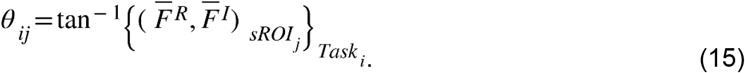

The resulting *F*_*WW*_ follows the *F*(*1, n-2*) distribution approximately, from which a *P*-value is derived and subsequently adjusted with a Bonferroni correction for three comparisons. For each task, only those sROIs with significant activations (surge height > 2.0) were included in the comparisons. The resulting numbers of sROIs were different across tasks (Supplementary Table 1e). The Watson Williams test was performed using the CircStat toolbox^82^ in Matlab (https://www.mathworks.com/matlabcentral/fileexchange/10676-circular-statistics-toolbox-directional-statistics).

### Traveling wave movies

To visualize the dynamic propagation of brain activity, we created an isophase traveling wave movie for each task using the *phasemovie*.*tcl* script in *csurf tksurfer* (Supplementary Movies 1-6). These animations resemble the motion of rainbands seen in a dynamic weather radar map, providing an intuitive visualization of spatiotemporal activations^41^. Each movie consists of 41 sequential frames between 4 and 12 s, each of which displays brain regions (as wavefronts) whose activation phases fall within a rotating sector spanning 9° (0.4 s) and advancing at a step of 4.5° (0.2 s).

## Acknowledgements

This research was supported by the University of Macau Development Foundation (EXT-UMDF-014-2021); University of Macau (MYRG-CRG2024-00047-ICI, MYRG-GRG2023-00239-FAH, CPG2023-00016-FAH, MYRG2022-00265-ICI, MYRG2022-00200-FAH, CRG2021-00001-ICI, CRG2020-00001-ICI, SRG2019-00189-ICI, PIDDA 2020, PIDDA 2019); Macau Science and Technology Development Fund (FDCT 0001/2019/ASE); National Institute of Health (R01 MH081990 to M.I.S and R.-S.H).

## Data availability

All data generated for this study are available in the main manuscript and supplementary files.

## Code availability

Custom codes for analyzing phase-encoded fMRI data and traveling waves are included in *csurf* (a FreeSurfer-compatible package) available at                                                                https://pages.ucsd.edu/~msereno/csurf/

