## Supplementary Materials for "Neural ensembles for music production recruit more language instruments as rhythmic complexity increases"

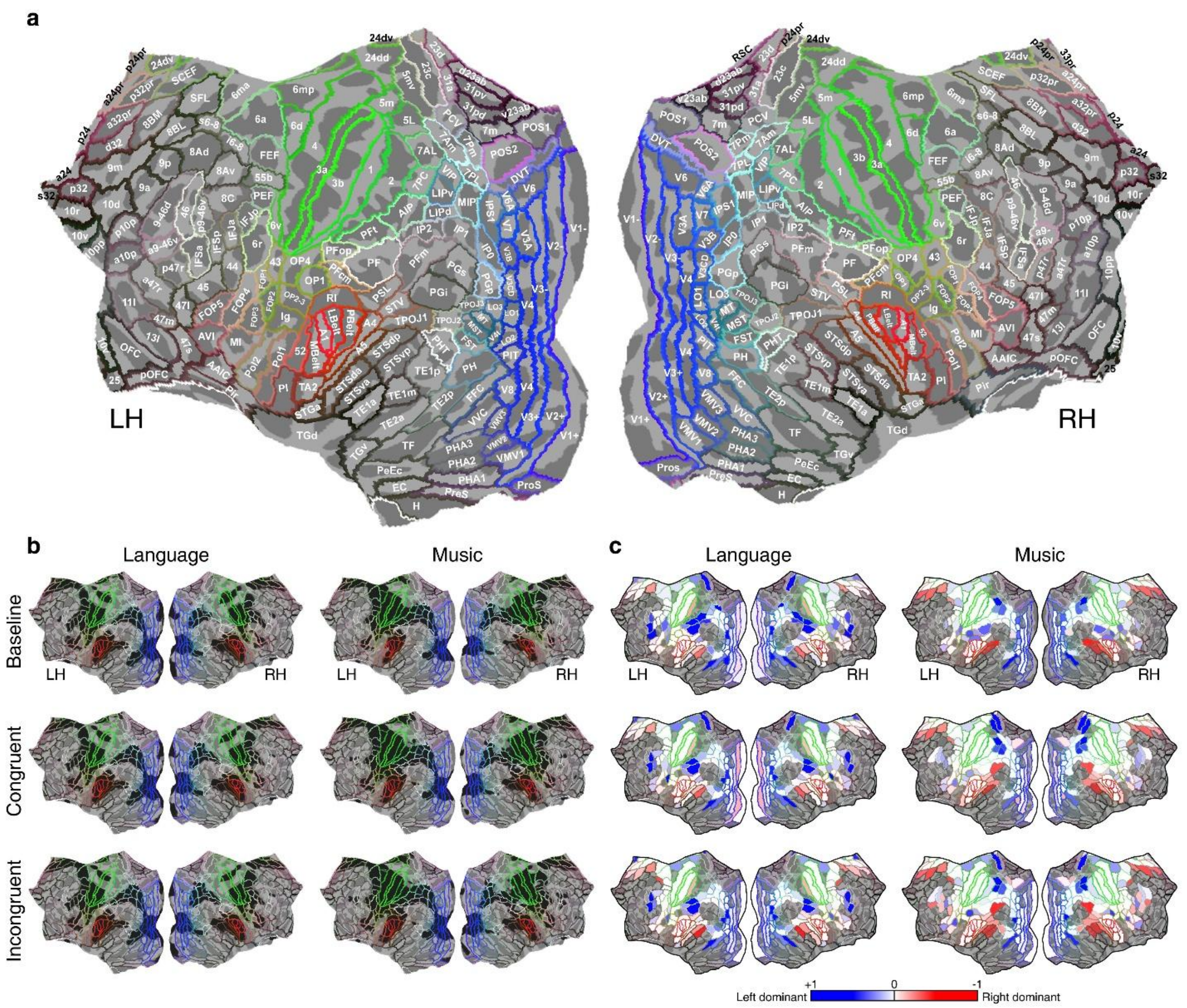


**Supplementary Figure 1. Surface-based regions of interest based on HCP-MMP1 parcellation. a,** HCP-MMP1 parcellation and sROI labels displayed on flattened cortical surface of *fsaverage*. **b,** Black regions indicate the extent of phase-encoded activations in Fig. 3a (columns 2 and 3), overlaid with HCP-MMP1 parcellation. **c,** Maps of laterality index (LI). Paired sROIs across hemispheres are shown in the same color (blue indicates left dominance; red indicates right dominance).

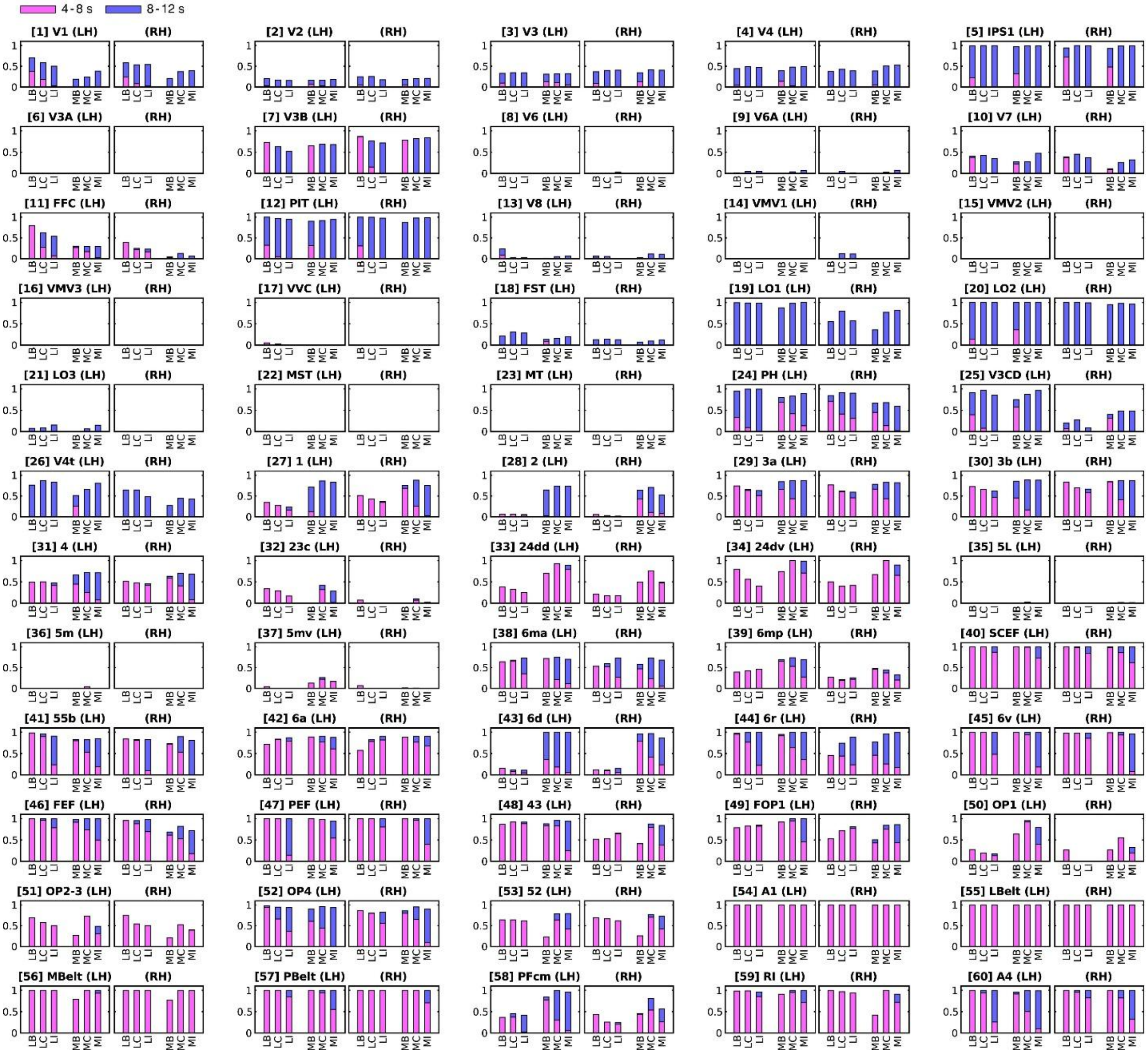


**Supplementary Figure 2. Activation ratios within surface-based regions of interest (No. 1–60).** The number in the square brackets indicates the sROI index in Supplementary Table 1b. Pink and blue segments represent the ratios of vertices with earlier-phase (4-8 s) and later-phase (8-12 s) activations, respectively. LB, LC, and LI: baseline, congruent, and incongruent conditions in language tasks; MB, MC, and MI: baseline, congruent, and incongruent conditions in music tasks. (LH): left hemisphere; (RH): right hemisphere.

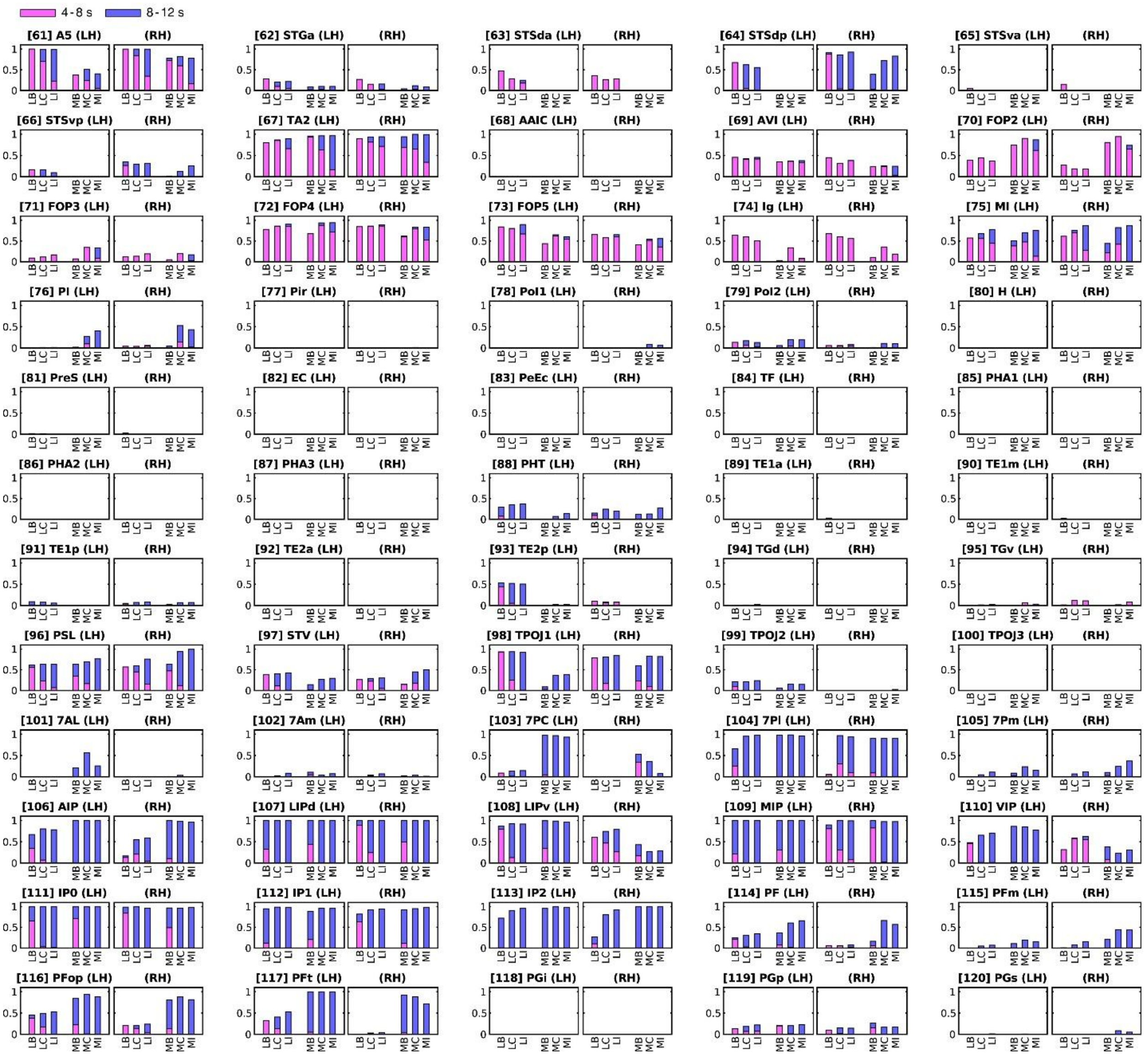


**Supplementary Figure 3. Activation ratios within surface-based regions of interest (No. 61–120).** All conventions follow Supplementary Fig. 2.

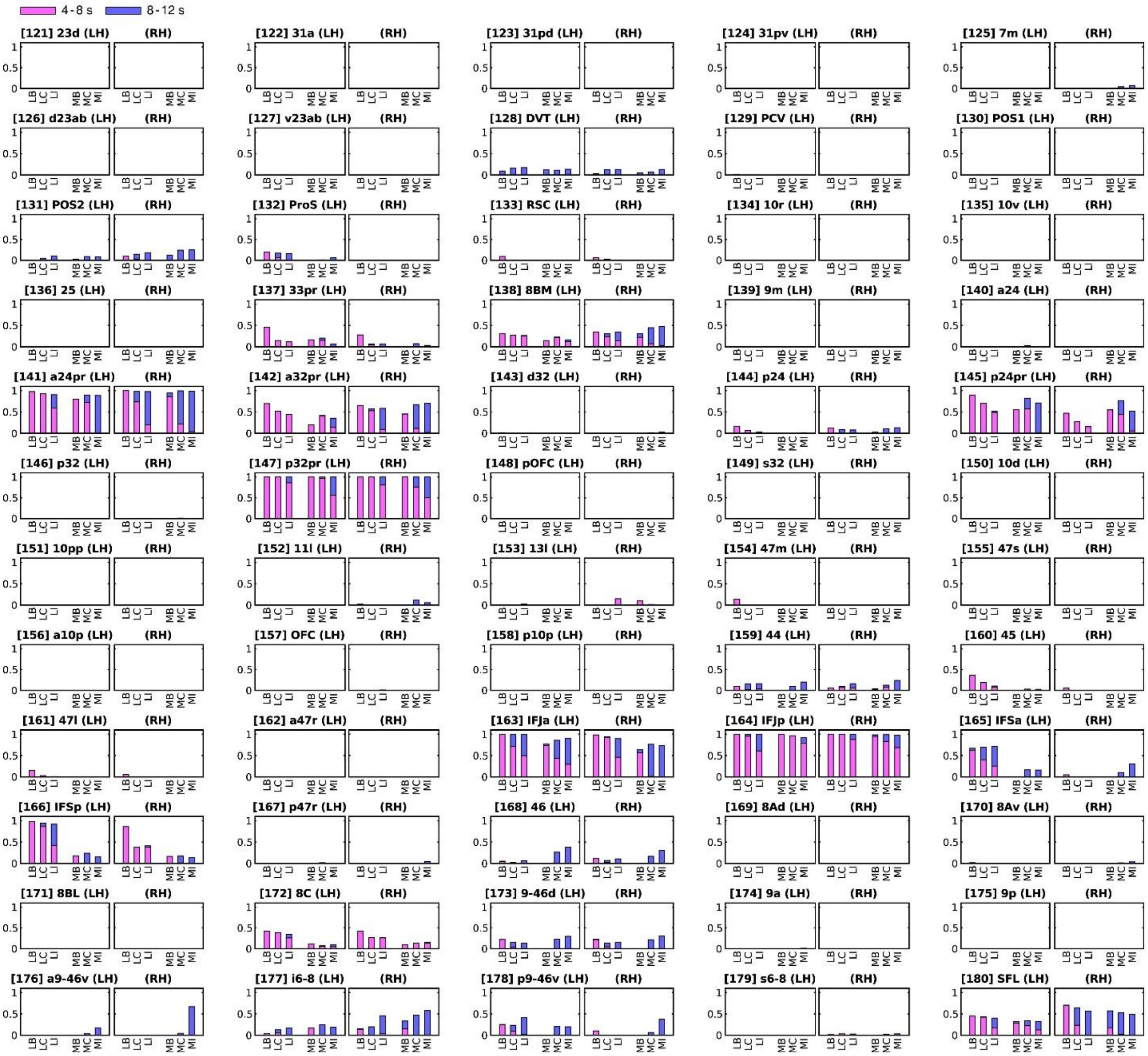


**Supplementary Figure 4. Activation ratios within surface-based regions of interest (No. 121–180).** All conventions follow Supplementary Fig. 2.

**Supplementary Table 1a to e | Detailed information for 180 sROIs based on the HCP-MMP1 atlas. a,** Activation ratios in 180 sROIs with activation phases between 4 and 12 s. **b,** Activation ratios in 180 sROIs for earlier-phase (4 - 8 s) and later-phase (8 - 12 s) activations. **c,** Percentage of sROI conjunction between language and music tasks for each condition. **d,** Laterality index (LI) for each sROI in each task. **e,** sROI mean phases. **Gantt chart (1)-(12).** Timing of above-threshold portions of surge profiles (height > 2.0; $F(2,230) = 4.7$, $P < 0.01$) in the Gantt charts in Fig. 5a,b, lower panels.

**Supplementary Movie 1. Animated Gantt charts and traveling waves of activations in language baseline condition.** Hemodynamic traveling waves visualize spatiotemporal information flows during the task. A vertical slider (spanning 180 sROIs) moves along the Gantt chart timeline (Fig. 5), highlighting sROIs with significant activations that correspond to spatial patterns on the flattened cortical surface. As the slider advances, these activation patterns form traveling waves that transition smoothly between brain regions over time. LH: left hemisphere; RH: right hemisphere.

**Supplementary Movie 2. Animated Gantt charts and traveling waves of activations in the language congruent condition.** All conventions follow Supplementary Movie 1.

**Supplementary Movie 3. Animated Gantt charts and traveling waves of activations in the language incongruent condition.** All conventions follow Supplementary Movie 1.

**Supplementary Movie 4. Animated Gantt charts and traveling waves of activations in the music baseline condition.** All conventions follow Supplementary Movie 1.

**Supplementary Movie 5. Animated Gantt charts and traveling waves of activations in the music congruent condition.** All conventions follow Supplementary Movie 1.

**Supplementary Movie 6. Animated Gantt charts and traveling waves of activations in the music incongruent condition.** All conventions follow Supplementary Movie 1.

**Supplementary Movie 7. Animated Gantt charts showing spatiotemporal changes across conditions of language and music tasks.** The movie illustrates rightward shifts of brain activation patterns as the Gantt charts change across three conditions of language and music tasks. LH, left hemisphere; RH, right hemisphere.
